# Immune cell senescence drives responsiveness to immunotherapy in melanoma

**DOI:** 10.1101/2025.07.31.667878

**Authors:** Pavlos Pantelis, Dimitrios Christos Tremoulis, Konstantinos Evangelou, Panagiotis Bakouros, Sophia Magkouta, Dimitris Veroutis, Orestis Ntintas, George Theocharous, Ioannis V. Kostopoulos, Eftychia Chatziioannou, Ioanna A. Anastasiou, Nefeli Lagopati, Dimitrios Skaltsas, Dimitris Kletsas, Dimitris Thanos, Alexandros J. Stratigos, Martin Röcken, Lukas Flatz, Russell Petty, George P Chrousos, Dimitris Vlachakis, Ourania E. Tsitsilonis, Timokratis Karamitros, Vassilis G. Gorgoulis

## Abstract

**Background:** Immunotherapy has significantly improved cancer treatment. However, it is not effective in all cancer patients, rendering the need to further delineate the differences among responders and non-responders at the molecular and cellular level. Unresponsiveness to immunotherapy has been attributed to dysfunctional immune cell states such as T-cell exhaustion and anergy, whereas the contribution of cellular senescence remains elusive. Herein, we have investigated the role of immune cell senescence in the response to checkpoint inhibitors in melanomas where these immunotherapies are applied as a first line treatment.

**Methods:** Two senescence detecting complementary approaches were utilized in a case control study we conducted. First, we implemented a senescence molecular signature we developed, termed “SeneVick” retrospectively in a single cell RNA-seq dataset from melanoma patients who received immunotherapy. Prior to this analysis, the signature was extensively validated in a variety of cell/tissue contexts, senescence types and species. Second, cellular senescence was assessed via an established experimental algorithmic approach in circulating immune cells of an analogous melanoma clinical cohort.

**Results:** Melanoma patients who did not respond to immunotherapy exhibited increased cellular senescence in their CD8+ T-cells, CD4+ T-cells, B-cells and NK cells compared to responders. This phenomenon was independent of patients’ age and not an outcome of immunotherapy, in contrast to conventional anti-cancer treatments. Interestingly, alterations of cell-cell interactions among the immune sub-populations in non-responders compared to responders were identified, suggesting the involvement of immune cell senescence in defective immune responses and treatment failure.

**Conclusion:** Overall, our findings support cellular senescence of the immune cell compartment within the TME, as a potent determinant of the response to immunotherapy and pave the way for strategies targeting immune cell senescence, as promising approaches to improve the outcome of such interventions.

## 1. Background

Following its initial discovery by Hayflick and Moorhead more than 60 years ago, as “aging at the cellular level”, noteworthy advancement has been achieved towards characterizing a cellular stress response mechanism that is distinct to the aging process, termed cellular senescence^1^. Physiologically and on a transient basis, senescence acts as a homeostatic mechanism, limiting the propagation of damaged cells in tissues. In contrast, if senescent cells are not timely eliminated by the immune system, they persist and accumulate, resulting in detrimental outcomes such as age-related pathologies and aging^1^.

For many years, a major drawback in the senescence field was the absence of reliable markers to effectively recognize senescent cells^2^. Identification of senescence relied mainly on the SA-b-Gal method, which is applicable only in cell culture and prone to false outcomes^3,4^. Moreover, other indirect and nonspecific for senescence markers were commonly applied. Overall, these approaches often resulted in misleading conclusions^3,5^. In order to bypass these obstacles, we and others recently proposed a senescence detecting algorithm (SDA) that increases the sensitivity and the specificity of senescence identification. It also allows for its accurate verification in any kind of biological sample, including formalin fixed and paraffin embedded (archival) material^1,3,4^. An essential component of SDA is the detection of lipofuscin, a hallmark and a common denominator of all senescent cells^1,3–5^.

The implementation of SDA retrospectively in clinical samples unveiled that cellular senescence is implicated in various pathologies such as cancer, COVID-19 disease, and giant cell arteritis, denoting that its role in the pathophysiology of human diseases largely remains encrypted and overlooked^1,6–8^. Interestingly, in cancer, one of the most common age-related diseases, it has been demonstrated that senescent cells act as a source of tumor recurrence via the senescence associated secretory phenotype (SASP) and/or the “escape from senescence” phenomenon ^1,4,9,10^, suggesting its involvement in the clinical outcome of cancer patients^11^. However, implementation of SDA in retrospective analyses will take time to provide results as it is labour-intensive, and in many cases the material is limited or even exhausted. Moreover, given the complex and largely heterogeneous nature of the senescence phenotype, tools that facilitate towards precise senescence identification are necessary to further elucidate its role in the pathophysiology and clinical course of human pathologies, such as cancer^12,13^. These conundrums and the unexplored for senescence abundant scRNA available databases led us to develop a molecular senescence signature, from now on termed “SeneVick” that could complement SDA in identifying senescence accurately^14^. As demonstrated, following extensive validation, SeneVick proved a highly efficient tool in demarcating non senescence from the senescence state.

Immunotherapy exemplified by checkpoint inhibitors has drastically influenced cancer therapy in the last decades^15^. These treatments aim to increase the efficacy of immune cells against the tumor. However, current cancer immunotherapies are not effective in all patients^16^. Cancer induced immunodeficiency is an important determinant of the response to such interventions, the molecular mechanisms though governing these processes remain to a large extent, unresolved^17^. Thus, an interesting matter that emerges regards the biological events that distinguish Responders (Rs) from Non Responders (NRs), to immunotherapy. Dysfunction of the immune cell compartment within the tumor microenvironment (TME), as an outcome of immune cell exhaustion or anergy, has been proposed while the involvement of immune cell senescence remains uncharted^5^.

Herein, we address this topic by implementing in a complementary manner experimental and *in silico* approaches, demonstrating that NRs to immunotherapy melanoma patients exhibit increased immune cell senescence in CD4+ and CD8+ T-cell, B-cell and natural killer (NK) cell populations compared to Rs.

## 2. Materials and Methods

### 2.1 Experimental planning

Prior addressing the main question of our study whether immune cell senescence drives responsiveness to immunotherapy in melanoma patients (Sections 2.7.1 and 2.7.2) we extensively validated our molecular senescence signature SeneVick in *in silico* (Sections 2.2.1, 2.3, 2.6.2) and experimental senescence models (Sections 2.2.2, 2.4, 2.5, 2.6.1).

### 2.2 Senescence Models

#### 2.2.1 *In silico* Setting

The analysis for the senescence control datasets which contained scRNA data from mice of different age groups and human fibroblasts (WI-38)^18,19^ was conducted as outlined below: The scRNA-seq data (GSE132042 and GSE226225) for the mice cohort and human fibroblasts (WI-38), respectively, were downloaded from GEO. Particularly, the first control dataset^19^ (GSE132042) contained scRNA-seq data from various tissues of mice belonging to six age groups, from 1 month to 30 months. The second dataset includes scRNA-seq data from human fibroblasts undergoing radiation- or therapy-induced senescence following Etoposide (ETO) treatment^18^. Cells with less than 1000 detected genes were omitted from the analysis. The gene counts were decontaminated using DecontX ^20^(v1.0.0). Seurat (v5.0.1) was used for the main part of the analysis. The quality control steps included filtering of cells that had a mean expression > 2^2.5-1 of selected housekeeping genes (**Table S1**) and the mitochondrial counts were removed. The cells were integrated with the fastMNN function (only in the fibroblast dataset) and clustered using a clustering resolution of 0.3 and 40 principal components.

In the mouse senescence control dataset, which contained a publicly available single-cell transcriptomic atlas that was extracted across the lifespan of *Mus musculus*^19^ linear models were used to determine whether there is a linear relationship between the timepoints and the senescence signature score. A Wilcoxon test was conducted per cell type between those two groups, in order to determine senescence score differences.

The senescence enrichment score of each signature^14,21,22^ (SeneVick, SenMayo and Fridman) in the human fibroblasts originating from the human senescent control dataset^18^ (GSE226225) was compared via Wilcoxon Test. Furthermore, we compared the enrichment levels of the aforementioned signatures in the human fibroblasts, in which the induction of senescence was accomplished with different senescent inducers, (irradiation, ETO treatment) and significance was also assessed via Wilcoxon Test. Segmented linear regression (segmented R package, v.2.1.3) was used in order to model the increase of senescence enrichment scores of the different senescence signatures across several timepoints, following ETO treatment.

#### 2.2.2 *In vitro* setting

Human diploid WI-38 fibroblasts (purchased by ATC, CCL 75) and Primary Human Fibroblasts (kindly provided by National Centre for Scientific Research Demokritos) of the three different age groups (7 years, 33 years and 75 years) were cultured in Dulbecco’s modified Eagle’s medium (DMEM, Biowest, L0104) supplemented with 10% FBS and 1% antibiotics. Cell cultures were maintained in an incubator at 37° C and 5% CO2. For etoposide (ETO) induced senescence, human diploid WI-38 fibroblasts were treated with 50 μM ETO (for six days, then cultured in regular medium without ETO-containing medium for four additional days. In the time course experiments, cells were collected at 0 (untreated), 1, 2, 4, 7, and 10 days after ETO treatment.

### 2.3 Analysis of the senescence signature SeneVick

#### 2.3.1 Gene Ontology-Based Functional Annotation

Gene Ontology (GO) enrichment analysis was performed to identify significantly overrepresented biological themes in SeneVick among its genes. GO is a hierarchically structured vocabulary encompassing three domains: Biological Process (BP), Molecular Function (MF), and Cellular Component (CC). Each domain captures different facets of gene function, allowing comprehensive annotation across cellular contexts. GO annotations and enrichment testing were conducted using g: Profiler^23^ and DAVID v6.8^24^ using the DAVID Knowlegebase v2023q4 as updated quarterly. Analyses were carried out using the Homo sapiens reference background (Ensembl GRCh38), and significance was determined using a hypergeometric test followed by Benjamini– Hochberg false discovery rate (FDR) correction. Only GO terms with FDR-adjusted p-values<0.05 were statistically significant. To complement the GO-based analysis, pathway annotations were retrieved from KEGG, Reactome, and BioCartadatabases. These included canonical signaling pathways relevant to senescence, such as the p53/p21 axis, NF-κB activation, mitochondrial metabolism, SASP regulation, and DNA damage response. Each gene was mapped to one or more functional categories based on GO and pathway term enrichment.

#### 2.3.2 Construction of Functional Association Matrix

A binary gene-function matrix was constructed in which rows represented individual genes and columns represented significantly enriched GO terms and pathways. Each matrix entry was coded as “1” if the gene was associated with a given term, and “0” otherwise. To reduce dimensionality and remove redundancy due to overlapping terms, Principal Component Analysis (PCA) was applied, retaining components explaining >90% of the total variance. To delineate distinct biological modules within the senescence signature, unsupervised clustering was performed on the reduced gene-function matrix. Two complementary approaches were applied:

1. K-means clustering^25^ was implemented using the Euclidean distance metric. The optimal number of clusters (k) was selected by evaluating the elbow plot and silhouette score across a range of k values. This method identified non-overlapping clusters of genes sharing similar functional annotation profiles.
2. Agglomerative hierarchical clustering was performed using Ward’s linkage method, producing a dendrogram to evaluate hierarchical relationships among functional groups. Final clusters were defined by cutting the dendrogram at a level that maximized within-cluster similarity while preserving between-cluster separation.

Genes associated with multiple terms were assigned to the cluster in which they showed the highest cumulative enrichment score. In cases of ambiguity, gene membership was resolved based on semantic similarity scoring, calculated using the GOSemSim R package. This enabled biologically meaningful classification based on ontological proximity to core senescence processes. All analyses were conducted in R version 4.3.0 and Python 3.10, using the packages clusterProfiler, factoextra, GOSemSim, scikit-learn, and SciPy. GO and pathway databases were accessed in June 2025 to ensure current annotation status.

### 2.4 Senescence assessment

Senescence assessment was performed in cells following double staining with the senescence detecting reagent GLF16 that we generated along with related senescence markers, and in tissues by applying GL13^26^ and GLF16, according to the SDA^3,27,28^, as follows:

#### 2.4.1 GLF16 staining/Immunofluorescence

Primary human skin fibroblasts of three different age groups (7 years, 35 years and 75 years) or human diploid WI-38 fibroblasts were seeded (2x10^5^ cells/well) on coverslips (12-mm diameter). The latter were subsequently treated with Etoposide for senescence induction^18^. In both cases, coverslips were subsequently removed; cells were washed fixed (4% PFA/PBS, 10min, 4°C) and permeabilized (Triton 0.3%/PBS 15 min). Blocking of non-specific epitopes was performed using sheep serum (dilution 1/40, S22, Merck Millipore). Cells were subsequently stained for lipofuscin using GLF16 for 10 min (70 mg/ml) avoiding light exposure as previously described^27^.Coverslips were washed 3 times for 10 min each with GLF16 diluent (2.5% DMSO/2.5% Tween-20/PBS). Then, cells were incubated with anti-p16^ΙΝΚ4Α^ (16D5, QR Labs) or -p21^WAF1/CIP1^ (1947S, Cell Signaling) antibodies for 1h in RT, followed by application of appropriate secondary antibodies (for 1h in RT). Nuclei were finally visualized by DAPI. Cells were washed (30s with dH2O) and coverslips were mounted onto slides for microscopy

In the case of tissues 4-μm-thick sections of formalin fixed and paraffin embedded tissues (FFPE) were obtained, de-paraffinized and hydrated. Antigen retrieval was performed by emerging samples in 10 mM of citric acid (pH 6.0) in a steamer for 15 min. Tissue samples were cooled down and washed with PBS. Blocking of non-specific binding for the epitopes was done by applying normal goat serum for 1 h at room temperature (dilution 1:40, Abcam, Cambridge, UK ab138478). The samples were incubated with the following primary antibodies overnight at 41C: CD4 (Ready to use, M7310, Dako), CD8 (1:20, 144B, Dako) and CD20 (1:150, 250586, Abbiotech). Positive cells were visualized using secondary goat anti-mouse (Abcam, ab6785, polyclonal) and goat anti-rabbit IgG H&L antibody (Alexa Fluor 488; 1:500; Abcam, ab150077, polyclonal) for 11h. Upon staining with primary and secondary antibodies, tissue sections were stained for lipofuscin applying GLF16 for 101min (601μg/ml) in the dark. Excess compound was removed by washing three times with the GLF16 diluent (2.5% DMSO/2.5% Tween 20/PBS). Nuclei were finally visualized by DAPI staining. The samples were washed (301s with dH2O), and coverslips were mounted onto slides for microscopy. Samples were imaged on a Leica TCS-SP8 confocal microscope.

#### 2.4.2 GL13 Staining

FFPE sections from melanomas were deparaffinized and hydrated. Subsequently, antigen retrieval was carried out as described in section 2.4.1 and after blocking of non-specific binding sites, the tissues were incubated sequentially in 50% and 70% ethanol for 5 min each, respectively. Following application of GL13 on each tissue, the samples were incubated at 37 °C for 10 min. At the end of this step, the samples were washed with 50% ethanol for 2-3 min, with PBS and then Triton-X 0.3%/PBS was applied for 5 min in order to remove any reagent precipitates. Tissues were washed again with PBS and anti-biotin antibody (Cat. no: K5007, in dilution 1:300, Hyb-8, ab201341, Abcam, Cambridge, UK) was applied and incubated for 1 h at RT. The mean percentage of GL13-positive cells was assessed from ≥5 high-power fields (Objection 40×) per sample using a ZEISS Axiolab5 optical microscope.

#### 2.4.3 Immunocytochemistry - Immunohistochemistry

Cells from each cell line were seeded on coverslips as mentioned above. For the Immunocytochemistry (ICC), the cells were permeabilized using Triton-X 0.3%/PBS for 15 min at RT, followed by the blocking of non-specific binding sites with goat serum ^29^ (Abcam ab138478, in 1:40) for 1 h at RT. Cells were then incubated with Ki67 (Cat.no: ab16667, dilution 1:250, SP-6, Abcam, Cambridge, UK) for 1 h at RT. Positive cells were visualized using the Dako REAL EnVision Detection System kit (Cat.no: K5007, Santa Clara, CA, USA) according to the manufacturer’s instructions using 3,31-Diaminobenzidine (DAB) (brown color). Coverslips were counterstained with hematoxylin, sealed and observed under a ZEISS Axiolab5 (Munich, Germany) optical microscope with 20× or 40× objectives.

Regarding the FFPE material (see section 2.4.1) sections were incubated with anti-CD4, anti-CD8 and anti-CD20 antibodies, respectively) overnight at 41C: CD4 (Ready to use, M7310, Dako), CD8 (1:20, 144B, Dako) and CD20 (1:150, 250586, Abbiotech). Positive cells were visualized using the Dako REAL EnVision Detection System kit (Cat.no: K5007, Santa Clara, CA, USA) according to the manufacturer’s instructions using 3,31-Diaminobenzidine (DAB) (brown color). Sections were counter-stained with hematoxylin and observed using a ZEISS Axiolab5 optical microscope with a 20×objective, 25µm scale bar).

### 2.5 Telomere Analysis

#### 2.5.1 Telomere Length Measurement

Relative telomere length determination (T/S) refers to the ratio of telomere (T) hexamer repeat sequence TTAGGG signal, to autosomal single copy gene (S) signal. To assess this, cells were collected and frozen at −80 °C until all samples were ready for simultaneous DNA extraction and analysis. Genomic DNA was extracted using the T3010 Monarch Spin gDNA Extraction Kit (T3010, New England Biolabs). Telomere (“T”) and single copy gene (human albumin, “S”) lengths were measured via real-time PCR (Roche LC480, Roche Diagnostics Corporation, Indianapolis, IN). Samples were loaded on 96-well plates and run in triplicate. Repeated measures of the T/S ratio in the same DNA sample gave the lowest variability when the sample well position for T PCR on the first plate matched its well position for S PCR on the second plate. When one sample’s duplicate T/S values differed by greater than 7%, the sample was run a third time, and the two closest values were averaged to give the final result. This ratio was subsequently normalized by control DNA samples to yield relative standardized T/S ratios proportional to average telomere length. A 5-point standard curve [made of pooled reference DNA samples (100 to 6.25 ng/uL) and randomly located internal QC sample replicates (n=5), were utilized as calibrator samples, to guide analysis and indicate overall quality of assay performance. Additionally, a non telomeric control was added to random well locations to provide a unique fingerprint for each plate. The primers (100μΜ, Integrated DNA Technologies Coralville, IA) used were the following:

i. telomeric assay: TelG [5’-ACACTAAGGTTTGGGTTTGGGTTTGGGTTTGGGTTAGTGT-3’] TelC [3’-TGTTAGGTATCCCTATCCCTATCCCTATCCCTATCCCTAACA-5’]
ii. single-copy gene (Albumin) assay: AlbU [5’-CGGCGGCGGGCGGCGCGGGCTGGGCGGAAATGCTGCACAGAATCCTTG-3’] AlbD [5’-GCCCGGCCCGCCGCGCCCGTCCCGCCGGAAAAGCATGGTCGCCTGTT-3’] PCR was performed using 20uL reaction volumes consisting of: 10 uL of 2X Luna® Universal qPCR Master Mix (NEB, US), 7.0 uL of MBG Water, and 0.5 uL of 1 µM primers mix. Thermal cycling was performed on a LightCycler 480 (Roche) where PCR conditions were (i) T (telomeric) PCR: 95°C hold for 5 min, denature at 98°C for 15 s, anneal at 54°C for 2 min, with fluorescence data collection, 35 cycles and (ii) S (single-copy gene, Alb) PCR: 98°C hold for 5 min, denature at 98°C for 15 s, anneal at 58°C for 1 min, with fluorescence data collection, 43 cycles. Ct values of triplicates were averaged, if meeting a CV threshold of less than 2%. The telomere (T) concentration was divided by the albumin (Alb) concentration (S) to yield a raw T/S ratio. Raw T/S ratios were subsequently normalized by average internal QC calibrator samples within the same plate set.Z-scores were calculated to adjust RTL in case differences in dynamic range are introduced by systematic differences between batches.

#### 2.5.2 Telomeric peptide nucleic acid (PNA) FISH

Telomeric PNA-FISH was held according to the latest established protocols^30^. The primary skin human fibroblast cells of the three different age groups (7, 35 and 75 years) were cultured in a confluency of 60–80% and they were split at 48–72 hours before harvesting for metaphase chromosomes. Cell pellets were fixed with methanol and acetic acid and dropped on wet slides for overnight incubation. Cells were subsequently re-hydrated using PBS (15 min, RT) and subsequently incubated with RNase A (100 μg/μl, Merck KGaA, Darmstadt, Germany) for 1 h at 37°C. Chromosome preparations were fixed in 3.7% formaldehyde (2 min) and washed withTBS (twice, 5 min each). Chromosome preparations were digested with pepsin (1 mg/ml, in 10 mM HCl,pH 2) at 37°C for 10 min and then washed twice with TBS and finally dehydrated by serial incubations in 70, 85, and 96% cold ethanol and air-dried. Telomere-specific hybridizations were accomplished employing Cy3-labeled (TTAGGG)3 and FITC-labeled (CCCTAA)3 PNA probes (BioSynthesis, Lewisville, TX). After two consecutive washing steps, one in PBS and one in Wash solution (0.1 M Tris–HCl, 0.15 M NaCl, 0.08% Tween-20, pH 7.5) at 65°C for 5 min, the slides and the probe were both preheated at 37°C for 5 min and next 10 μl of hybridization mixture comprising of 0.2–0.8 μM PNA telomeric probes, 70% formamide, and 10 mM Tris, pH 7.2 (Cytocell, Oxford Gene Technology, UK), was positioned on the marked area of the slide. The latter underwent heating at 80°C for 5 minutes during denaturing FISH, whereas this step was excluded in the non-denaturing FISH protocol. Slides from both denaturing and non-denaturing FISH procedures were incubated overnight at 37°C in a humid environment. On the following day, slides were sequentially washed: once in PBS for 15 minutes, once in 0.5× Saline Sodium Citrate buffer (SSC) containing 0.1% SDS at 72°C for 2 minutes, once in 2× SSC (pH 7) supplemented with 0.05% Tween-20 at room temperature for 30 minutes, and twice more in PBS for 15 minutes each. Preparations were then counterstained and mounted with Vectashield containing DAPI (Vector Laboratories Inc). Images were captured under an Axion Imager Z1 Zeiss fluorescence microscope (63x objective) and analyzed using MetaSystems Isis software (Perner et al.2003). The signal of the centromere of chromosome 2 functioned as the internal reference control. Human centromere 2-specific PNA probes labeled with Cy3 or FITC were provided from DAKO Cytomation(Glostrup, Denmark).

### 2.6 Transcriptomics and analysis

#### 2.6.1 Transcriptomics

Primary human Fibroblasts of 7, 33 and 75 years old donors were seeded onto 10-cm cell culture plates (70% confluency). Cells were collected and total mRNA was extracted using the NucleoSpin RNA mini kit (Macherey-Nagel, Germany). RNASeq libraries were prepared with the NEBNext ultra II directional RNASeq kit (Reverse strand specificity) and single end sequenced at 101 bp length with the Illumina NovaSeq 6000 platform, in the Greek Genome Center of BRFAA.

Raw data were mapped to the human genome (version GRCh38/hg38) using STAR^31^ (aligner. Samtools^32^ were used for data filtering and file format conversion, while the HT-seq count algorithm (Anders et al. 2015) was used to assign aligned reads to exons using the following command line ‘‘htseq-count –s nom intersection –nonempty’’. Normalization of reads and removal of unwanted variation was performed with RUVseq^33^. Differentially expressed genes were assessed using the DESeq2 R package^34^ and the significant genes were characterized by log2 fold change cut-off of 0.5 and p value less than 0.05. Gene ontology and pathway analysis was accomplished using the Database for Annotation, Visualization and Integrated Discovery^35^ (DAVID). Only pathways and biological processes with p value less than 0.05 were characterized as significantly enriched. Heatmaps representing the significant differentially expressed genes and the most significant genes where SeneVick found enriched were constructed with R package Shiny^36^, where hierarchical clustering was performed, with linkage method ‘average’.

#### 2.6.2 Gene set Enrichment Analysis

Gene Set Enrichment Analysis^37^ (GSEA) was used in order to determine whether SeneVick is enriched in senescent samples. More specifically, the signature’s genes that are expected to be upregulated were tested for enrichment separately from those expected to be downregulated. Age in years was treated as a continuous variable and pearson correlation was used to determine the enrichment or depletion of the signature’s genes across age.

### 2.7 Melanoma patients

#### 2.7.1 *In silico* datasets

##### 2.7.1.1 Integration of databases

The scRNA-seq data^38,39^ (PMC6641984 and PMC6410377) for the melanoma cohort (**Table S2**) were processed as described above (Section 2.1), with the additional steps of a) the conversion of the counts to TPM, log transformation and b) the use of a minimum mean housekeeping expression threshold of log2(TPM+1) > 2.5. After the quality control steps, the cells were annotated using SingleR (v2.4.1) while using as a reference, a dataset comprising 300,000 immune cells^40,41^. The response status of the melanoma patients in the *in silico* datasets was assessed according to the Response Evaluation Criteria in Solid Tumours (RECIST)^42^.

The dataset was then split into broad cell types and comparisons were made between Rs and NRs. For each cell type, the dataset was integrated using the fastMNN function and clustered using a clustering resolution of 0.3 and 40 principal components. Ucell^43^ (v2.6.2) was used to calculate the senescence score, based on each senescence signature: SeneVick, SenMayo and Fridman^21,22^ and the T-cell anergy and exhaustion score. Next, in case a cell type contained clusters that exhibited low senescence score and high exhaustion score, this cluster was excluded. A Wilcoxon test was conducted between the Rs and NRs cells, in order to determine senescence score differences. The scRNA data obtained from patients prior to immunotherapy administration were analyzed using the same method. Of note the only difference was noted in the fact that there wasn’t need for integration of the data and that in cases where the low senescence - high exhaustion cells were not forming a separate cluster such as in CD4+ T-cells, we removed all the cells which had a senescence score below the 20th percentile and simultaneously an exhaustion score above the 80th% percentile and used a clustering resolution of 0.5.

##### 2.7.1.2 Cell Communication Analysis

In order to infer the cell-cell communication between the immune cells the CellChat R (v2.2.0) package was used^44^. The cells were grouped based on their cell type and the response status and the minimum number of cells required in each cell group for cell-cell communication was set to 50. Subsequently, the differences between Rs and NRs Ligand-Receptor pair communication probabilities in CD8+ T-cells, CD4+ T-cells, NK cells and plasma cells were identified.

#### 2.7.2 *Ex vivo* melanoma setting

Twenty-four (24) melanoma patients that received single or combinational immune checkpoint inhibitors, as first line therapy after entering stage IV, were analyzed. Patient and clinical characteristics are depicted in **Table S3**. All patients included in this study gave their written consent and the study was approved by the local ethical review board (project ID: Ethikkommission Ostschweiz, EKOS 16/079). Patient’s response was assessed using the Response Evaluation Criteria in Solid Tumours (RECIST) version 1.1^42^, approximately 3 months after initiation of therapy and at 3-month intervals thereafter. The Best Overall Response (BOR), defined as the best response recorded from the start treatment initiation until disease progression was documented. Based on BOR, patients were categorized as responders^42^ (complete or partial response) or nonresponders (NR) (stable disease or progressive disease).

##### 2.7.2.1 Senescence assessment in PBMCs

PBMCs from the melanoma patients were obtained following established procedures using Ficoll as previously described^45^. Peripheral blood mononuclear cells (PBMCs) were thawed, washed twice with PBS containing 0.5% heat-inactivated fetal bovine serum (PAN-Biotech GmbH; staining buffer), and resuspended in staining buffer. Cell viability and concentration were assessed microscopically using a hemocytometer (Neubauer chamber) and Trypan blue staining (Corning®, NY, USA). Cells were set at a final concentration of 5x106/ml and 100 μL were transferred to a 5 ml round-bottom polysterene FACS tube (BD Biosciences, NJ, USA). Cells were labelled with BD Horizon™ Fixable Viability Stain 570 (BD Biosciences, NJ, USA) for 20 minutes at 4°C in the dark and washed twice with staining buffer, before adding the master mix of 15 fluorochrome-conjugated monoclonal antibodies (**Table S4**) targeting surface antigens (BD Biosciences, NJ, USA; Biolegend Inc., CA, USA). To minimize non-specific binding, 10 μL of BD Pharmingen™ MonoBlock™ buffer (BD Biosciences, NJ, USA) was added together with the antibody master mix, and cells were incubated for 30 minutes at 4°C in the dark. Further, cells were fixed and permeabilized using the BD Pharmingen™ Human FoxP3 Buffer Set (BD Biosciences, NJ, USA), following the manufacturer’s instructions, and labelled with anti-Ki67 antibody (BD Horizon™ BV711 Mouse Anti-Human Ki67; BD Biosciences, NJ, USA) for 30 minutes at 4°C in the dark. After washing twice with staining buffer, cell pellets were resuspended in 200 μL GLF16 diluent [PBS, 2.5% Tween-20 (Sigma-Aldrich®), 2.5% DMSO (PAN-Biotech GmbH)] containing 2 μl of the GLF16 dye (200 μg/mL) and incubated for 10 minutes at room temperature in the dark under mild shaking. After two washing steps with GLF16 diluent, samples were acquired on a Cytek Northern Lights spectral flow cytometer (Cytek® Biosciences) For stable flow cytometer performance, daily SpectroFlo® QC Beads (Cytek® Biosciences) were run. Data analysis was performed with FlowJo™ v10 Software (BD Life Sciences). The gating strategy for analysis is presented in the supplemental material. During the aforementioned analysis unstained controls were analyzed for all samples.

##### 2.7.2.2 Senescence assessment in tissues

Senescence assessment in FFPE samples was carried out as described in Sections 2.4.1-2.4.3.

### 2.8 Quantification and Statistical Analysis

In each experiment, values are demonstrated as means ± standard deviation. Differences between groups were estimated using the parametric 2-tailed Student’s t test, Wilcoxon Test, the non-parametric Mann Whitney or 1-way ANOVA with Bonferroni’s post hoc test for multiple comparisons, as appropriate. p < 0.05 were considered significant. In order to compare the ages of Rs and NRs melanoma patients, a Shapiro-Wilk normality test was conducted to test the normality of the distributions of the age in each group (p<0.05, not normally distributed) and then a Wilcoxon test was held between the ages of the two groups of patients. Statistical analysis was performed using the Statistical Package for the Social Sciences v.13.0.0 (IMB).

## 3. Results

### 3.1 Decoding the senescence molecular signature SeneVick

We have recently generated Senevick by incorporating studies that assessed cellular senescence in human cells using the senescence detecting algorithm (SDA) and concurrently included a variety of high throughput data (transcriptomics: RNA-seq and scRNA-seq, proteomics and epigenomics^14^, **Figure 1a**). The signature is composed of 100 genes and exhibits an expression motif that complies with the senescence phenotype. The majority of them are down-regulated (n=67) (**Figure 1b**). Characteristically, the genes included in the signature are implicated in diverse biological processes and can be grouped into distinct functional clusters, reflecting to a large extent the hallmarks of the senescence phenotype^1^ (**Figure 1c**). The most profound cluster regards genes encoding potent cell cycle regulators and factors involved in the cellular response to DNA damage (**Figure 1c**). The second one consists of genes controlling chromosome structure and stability (**Figure 1c**). A considerable proportion of the signature encompasses genes related to the activation of immune responses and interactions among immune cells, while the other two clusters contain genes involved in metabolic and other functions (**Figure 1c**). Further zooming into clusters uncovered the involvement of the genes in a variety of cellular processes (**Figure S1a**). Interestingly, common genes between SeneVick and state of the art senescence molecular signatures namely the SenMayo and FridMan gene sets, were identified^21,22^ (**Figure S1b**).

**Figure 1:**
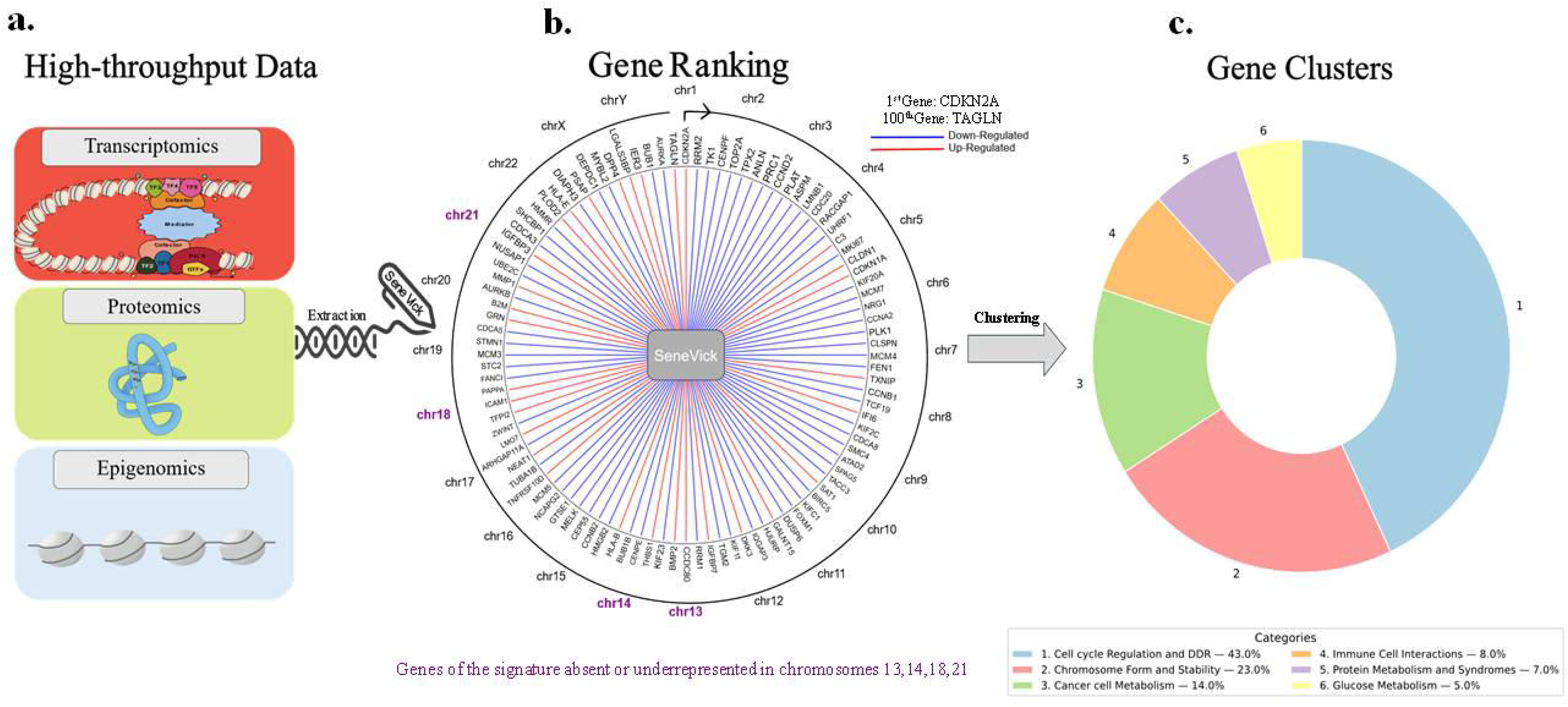
The senescence molecular signature, SeneVick. Pipeline followed for SeneVick extraction (a), SeneVick’s genes directionality and composition (b) and how these genes are clustered in a pie chart (c), is depicted. *The *CDKN2A* gene (p16^INK4A^) is commonly unattainable to be detected in high throughput data due to several technical factors. Bulk RNA-seq might not capture low-expressing transcripts and if p16^INK4A^ transcripts are degraded into fragments, short-read RNA-seq might fail to capture full-length sequences^74^.

Our senescence signature exhibits several unique and biologically intriguing features that underline its potential functional importance. First and foremost, analysis of the SeneVick gene set revealed that the constituent genes do not represent simple haplotypes, suggesting that their co-occurrence is not the result of genetic linkage or population-based inheritance patterns. Another prominent molecular feature of the SeneVick-encoded proteins is their strong enrichment in ankyrin repeat domains. Ankyrin repeats are highly structured motifs that mediate protein–protein interactions, often serving as scaffolds in large signaling complexes^46^. Their consistent appearance across nearly all proteins encoded by the SeneVick genes suggests a non-random, biologically meaningful pattern, potentially orchestrating the senescence phenotype. Indeed, ankyrin repeat-containing proteins have been implicated in the regulation of cellular integrity, cell-cycle arrest, stress signal transduction, and differentiation processes, all of which have been linked with cellular senescence^46^. In addition to structural motifs, the genomic architecture of the SeneVick genes revealed another layer of functional organization: a statistically significant enrichment in T-dimeric motifs within their genomic sequences, exhibiting a periodic distribution. Given that periodic nucleotide motifs have been associated with the dynamic modulation of gene expression, this feature raises the possibility of a regulatory role during senescence^47^. Lastly, when mapping the SeneVick genes on chromosomes, we identified their absence in chromosomes 14, 18 and 21 and their underrepresentation in chromosome 13 (**Figure 1b**). Interestingly, these chromosomes are associated with trisomy syndromes—Patau (trisomy 13), Edwards (trisomy 18), and Down syndrome (trisomy 21) or lethality (trisomy 14), entities characterized by premature aging, chronic inflammation and senescence phenotypes^48–50^. Thus, the absence or underrepresentation of SeneVick genes at these chromosomes may imply a protective genomic architecture, where senescence regulators are compartmentalized away from the chromosomal loci whose abnormal dosage leads to accelerated aging syndromes or premature death.

### 3.2 SeneVick effectively identifies senescence irrespective of tissue origin, senescence type or species

The SeneVick signature emerged by exploiting data from human cells of different tissue origin and proved efficient in demarcating non-senescence from senescence and in discriminating cellular senescence from aging in the liver^14^. We subsequently focused on testing its applicability and validating its fidelity in detecting senescence across tissues in other species besides humans. To elucidate this, we initially applied SeneVick in a publicly available single-cell transcriptomic atlas that was extracted across the lifespan of *Mus musculus*^19^. This dataset comprised single cell RNA sequencing (scRNA-seq) data obtained from 20 tissues and organs of mice split to six age groups, that ranged from 1 month which is equivalent of human early childhood to 30 months (equivalent of a human centenarian)^19^. Given that single cell analyses allow for the determination of gene expression in specific cell populations, they can facilitate uncovering certain cellular processes, such as cellular senescence, that might have been overlooked or hidden upon bulk RNA analyses. As demonstrated in **Figure S2a**, SeneVick was found significantly enriched in a variety of cell types and most profoundly in cardiac fibroblasts, keratinocytes and skeletal muscle cells of old mice (18 months and beyond, Wilcoxon, p<0.05), reaching the highest values in older mice (**Figure S2b**). These results are in line with other studies in the same tissues supporting a linear increase of the proportion of cells expressing senescence markers with age progression^19^. The latter was further confirmed in an *in vitro* human setting consisting of primary skin fibroblasts obtained from 7, 35 and 75 year-old individuals, capturing time-points of the aging process. In contrast to young cells, those from 75 year old donors were found to exert replicative senescence (a senescence type induced by telomere shortening)^1^ that was verified applying the SDA along with telomere length analyses (**Figure 2a, 2b, 2c, 2d, 2e**). To crosscheck this finding, we isolated RNA from these cells and implemented SeneVick in RNA seq data extracted from these fibroblasts and identified a progressive enrichment of the signature upon replicative senescence and age (**Figure 2f**). Particularly, GSEA analysis resulted in a positive Normalized Enrichment Score (NES) of 1.76 (p<0.002) for genes whose expression increases with age and a negative NES of -3.18 (p<0.001) for those that are downregulated as age progresses (**Figure 2f**). Overall, these findings highlight the potency of the extracted signature in detecting senescence, irrespective of tissue origin and species.

**Figure 2:**
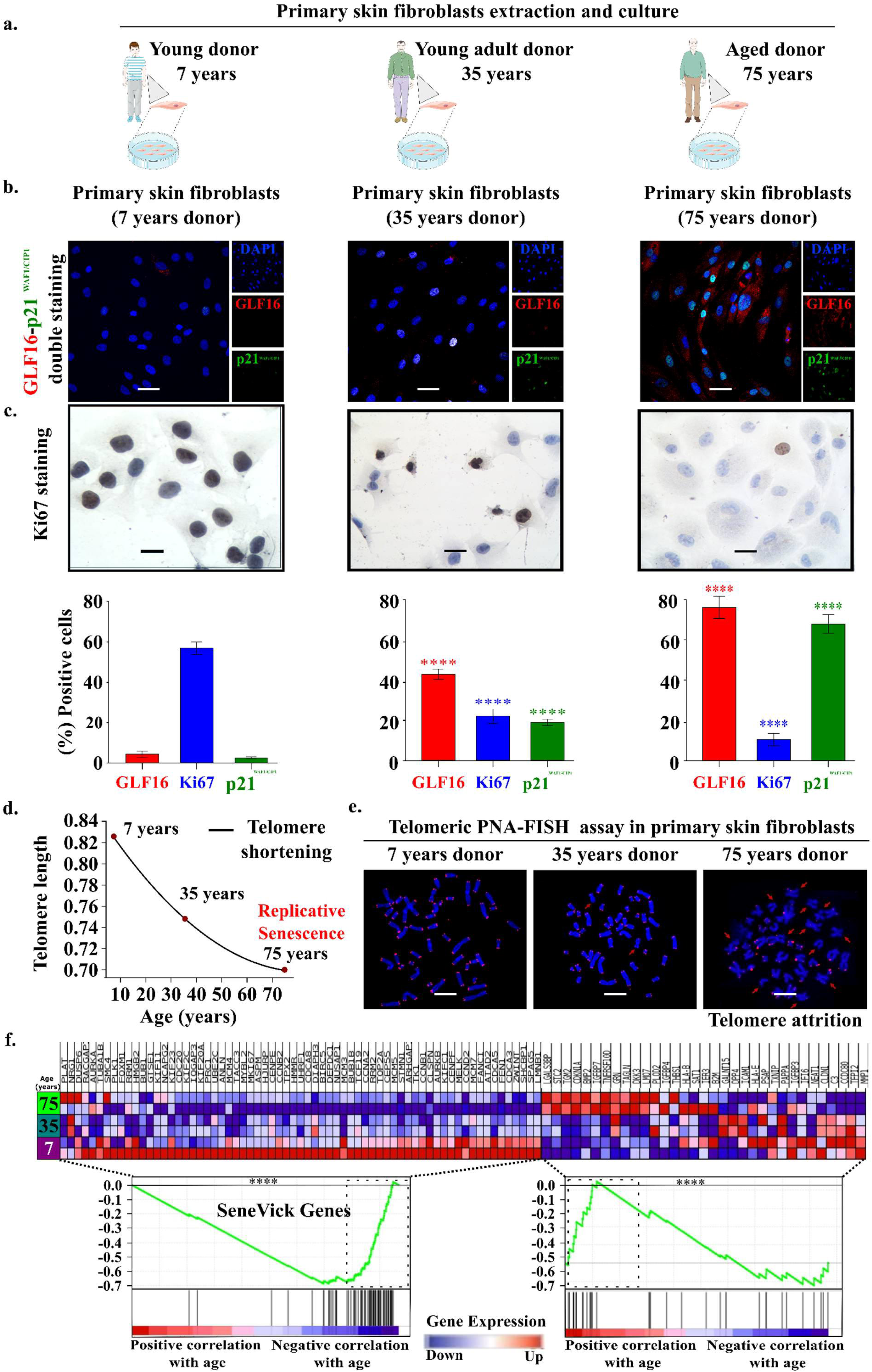
Validation of SeneVick in a human replicative senescence (aging) model. **a.** Schematic illustration of the primary skin fibroblasts extraction and culture **b.** Senescence assessment in human primary fibroblasts from different age groups (Age: 7, 33 and 75 years) using the senescence detecting algorithm (SDA). Representative images of double staining of cells with the senescence markers GLF16 (red) and p21^WAF1/CIP1^ (green) and DAPI counterstain. The images were quantified using ImageJ (*n*1=13 biological replicates). Objective: ×20. Scale bar: 301μm. **c.** Evaluation of proliferation in human primary fibroblasts from different age groups (Age: 7, 33 and 75 years). Representative images of Ki67 immunocytochemical staining (upper panel). Positive cells were calculated by evaluating the strong brown nuclear signal for Ki67. Graphs depict the percentage of positive cells (%) for GLF16, p21^WAF1/CIP1^ and Ki67 (lower panel). Approximately 100 cells per optical field were counted, and ≥5 high-power fields per sample were used for the quantification. Statistical analysis was performed employing unpaired t-test. The data obtained represent means ± standard deviation. *P*1<10.05, \*\**P*1<10.01, \*\*\**P*1<10.001, \*\*\*\**P*1<10.0001. Objective 20×, 40×. Scale bars: 30μm and 60 μm respectively. **d.** Telomere length curve depicting telomere attrition during aging in human primary fibroblasts. **e.** Microscopy images from PNA FISH using DAPI with a telomere-specific probe (right) depicting telomere attrition on metaphase chromosomes from the three age groups. Objectives 63x. **f. Left panel:** GSEA presenting the downregulated genes of SeneVick across three age groups of primary human fibroblasts (Age: 7, 33, and 75 years). A corresponding heatmap presents gene expression levels within these conditions. **Right panel:** GSEA demonstrating the upregulation of genes of SeneVick under identical conditions. The corresponding heatmap provides an in-depth visualization of these gene expressions levels. Two data sets were compared with unpaired t-tests, \**P*1<10.05, \*\**P*1<10.01, \*\*\**P*1<10.001, \*\*\*\**P*1<10.0001 “Donor icons were provided by Servier Medical Art (https://smart.servier.com/), licensed under CC BY 4.0 (https://creativecommons.org/licenses/by/4.0/).”

To further validate SeneVick, we compared it with two state of the art senescence signatures, namely the SenMayo and the Fridman^21,22^. Both signatures have been extracted from human data; SenMayo gene set consists predominately of senescence associated secretory phenotype (SASP) factors, while in the Fridman signature the key genes represented are involved in six particular pathways: pRB/p53, cytoskeletal formation, interferon-related, insulin growth factor-related, MAP kinase and oxidative stress. We implemented these three senescence signatures in a publicly available human sc-RNAseq dataset^18^ (GSE226225) obtained from human fibroblasts that were analyzed in two different settings. First, sc-RNAseq data were extracted from a time course experiment by monitoring cells for 10 days following treatment with the chemotherapeutic drug Etoposide (ETO) (**Figure 3a**). We repeated this experiment staining cells according to the SDA in three different timepoints^3^ (Day 0, 4, 10). Both approaches revealed absence of SeneVick enrichment and lack of senescence markers in day 0 and a progressive increase in the following days, reaching the highest values at day 10 (**Figure 3b, Figure 3c, Figure S3a**). The *in silico* dataset was used to compare SeneVick with the other two signatures taking into account the distribution of the enrichment scores and using segmented linear regression analysis (p<0,05), Senevick was not found enriched in non-senescent (control) cells (day 0) (**Figure S3b**). Similar observations emerged from the second setting where the signatures were applied in sc-RNAseq data of fibroblasts exerting different types of cellular senescence (irradiation-induced and ETO-induced) (**Figure S3c**). Altogether, SeneVick is more sensitive and efficient compared to the other signatures in demarcating senescent cells from non-senescent ones, even when senescence is low providing thus a valuable tool to uncover senescence that might be encrypted or overlooked. Of note, the senescence enrichment scores were calculated via UCell^43^, which takes into account which genes are up- and downregulated within the signature applied. However, this information is present only in the case of SeneVick and Fridman signatures, but not in the SenMayo one, providing a potential explanation of the cause of the relatively higher SenMayo enrichment score in the control group.

**Figure 3:**
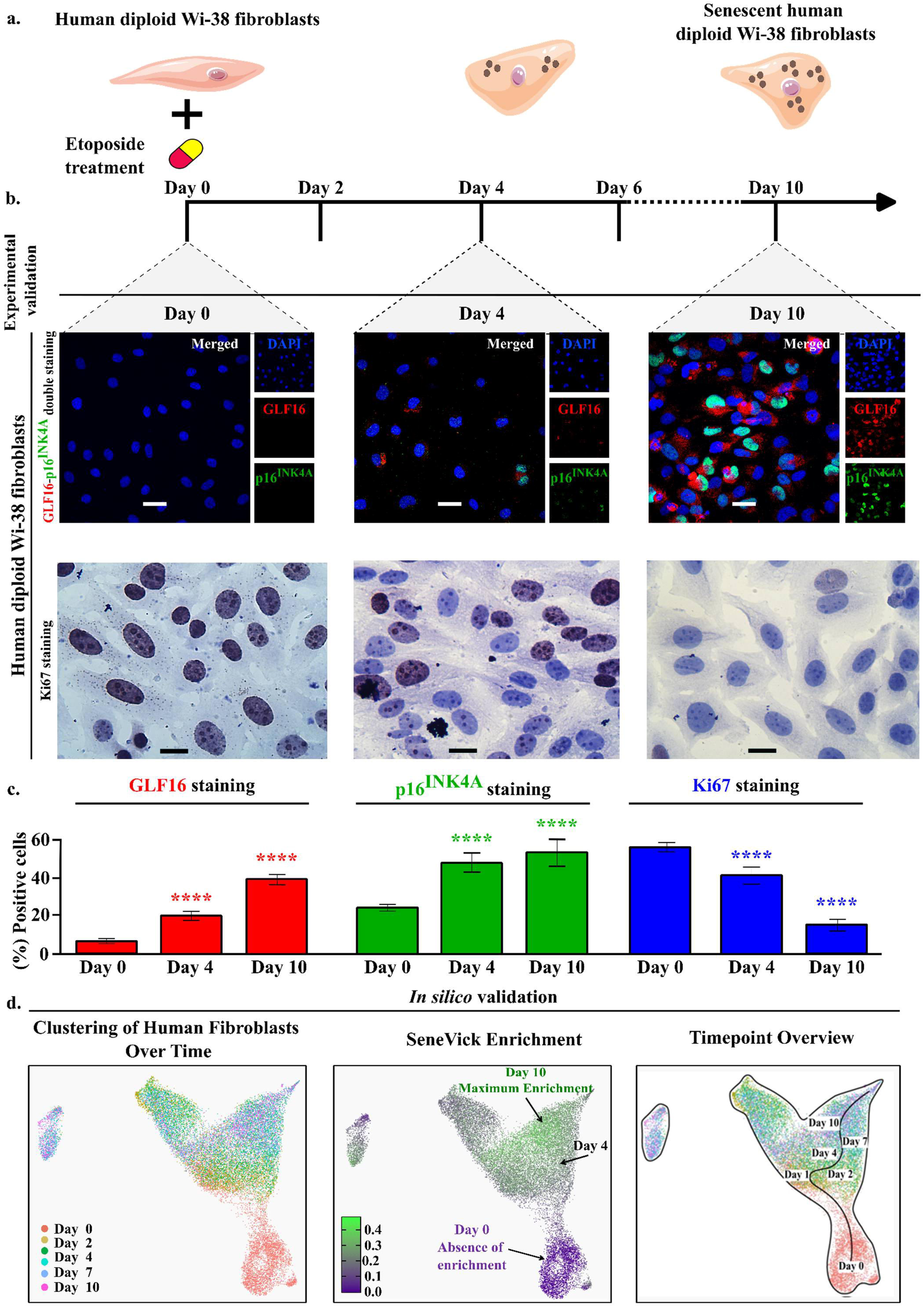
Validation of SeneVick in a human therapy induced senescence model. **a.** Schema depicting the experimental procedure followed to establish etoposide induced cellular senescence in human fibroblasts. **b.** Senescence assessment in human WI-38 fibroblasts using the senescence detecting algorithm (SDA). Representative images of double staining of cells with the senescence markers GLF16 (red) and p16^INK4A^ (green) and DAPI counterstain. The images were quantified using ImageJ (*n*1=13 biological replicates). Evaluation of proliferation in human WI-38 fibroblasts. Representative images of Ki67 immunocytochemical staining. Positive cells were calculated by evaluating the strong brown nuclear signal for Ki67. **c.** Graphs depict the percentage of positive cells (%) for GLF16, p16^INK4A^ and Ki67. Approximately 100 cells per optical field were counted, and ≥5 high-power fields per sample were used for the quantification. Statistical analysis was performed employing the Wilcoxon nonparametric test. The data obtained represent means ± standard deviation. * *P*1<10.05, \*\**P*1<10.01, \*\*\**P*1<10.001, \*\*\*\**P*1<10.0001. Objective 20×, 40×. Scale bars: 30μm and 60 μm respectively. **d.** Left panel: UMAP plot of human fibroblast scRNA data (GSE226225) displaying their clustering upon time (days), following etoposide treatment. Middle Panel: UMAP plot categorizing cells based on SeneVick enrichment upon time (days) in the GSE226225 dataset. Right panel: Indication of the different timepoints on UMAP plot of the scRNA data from human fibroblasts in the GSE226225 dataset. “Fibroblast cells icons were provided by Servier Medical Art (https://smart.servier.com/), licensed under CC BY 4.0 (https://creativecommons.org/licenses/by/4.0/).”

### 3.3 Immune cell senescence drives response to immunotherapy in melanoma patients

Next, we tested whether immune cell senescence contributes to dysfunction of the immune cell compartment within the TME, affecting the outcome of immunotherapy. Particularly, we conducted a case control study, implementing two senescence detecting approaches that complement each other. First, SeneVick was retrospectively applied in a single cell RNA-seq (scRNA-seq) dataset from melanoma patients that received immunotherapy^38,39^ (PMC6641984 and PMC6410377) and secondly, we performed the SDA in clinical material from an analogous melanoma cohort (**Tables S2 and S3**). We focused on melanoma based on the fact that immunotherapy is a first-line therapy in this type of malignancy, while in other types of cancer it is usually implemented in combination with chemotherapy or radiotherapy.

Regarding the first approach we took advantage of the only two identified in the literature studies containing single cell data from melanoma patients following immunotherapy, particularly PD1, CTLA4 or combined inhibition and concurrently demonstrating the response status of these patients^38,39^ (PMC6641984 and PMC6410377, **Table S2**). Thus, data regarding gene expression per cell type in Rs and NRs were accessible. In some cases, data prior treatment were also available (**Table S2**). As an initial step, we followed a detailed bioinformatic pipeline to integrate the two datasets as depicted in **Figure S4**, which included several steps of data processing, cell annotation and normalization. This process resulted in a cohort of a total of 48 melanoma patients comprising of 30 NRs and 18 Rs. In this setting, we found distinct clusters of the immune cell compartment comprising mainly of CD4+ T-cells, CD8+ T-cells, NK and plasma cells. During the ensuing stages, we implemented SeneVick in the latter dataset and investigated the senescence status in each immune cell population and in relation to the response outcome. As demonstrated in **Figure 4**, SeneVick enrichment was evident in CD4+ T-cells, CD8+ T-cells, NK and plasma cells and in relation to the response status we found that cells belonging to NR patients exhibited higher enrichment scores compared to responding patients. This phenomenon was independent of patient’s age (p=0.12).

**Figure 4:**
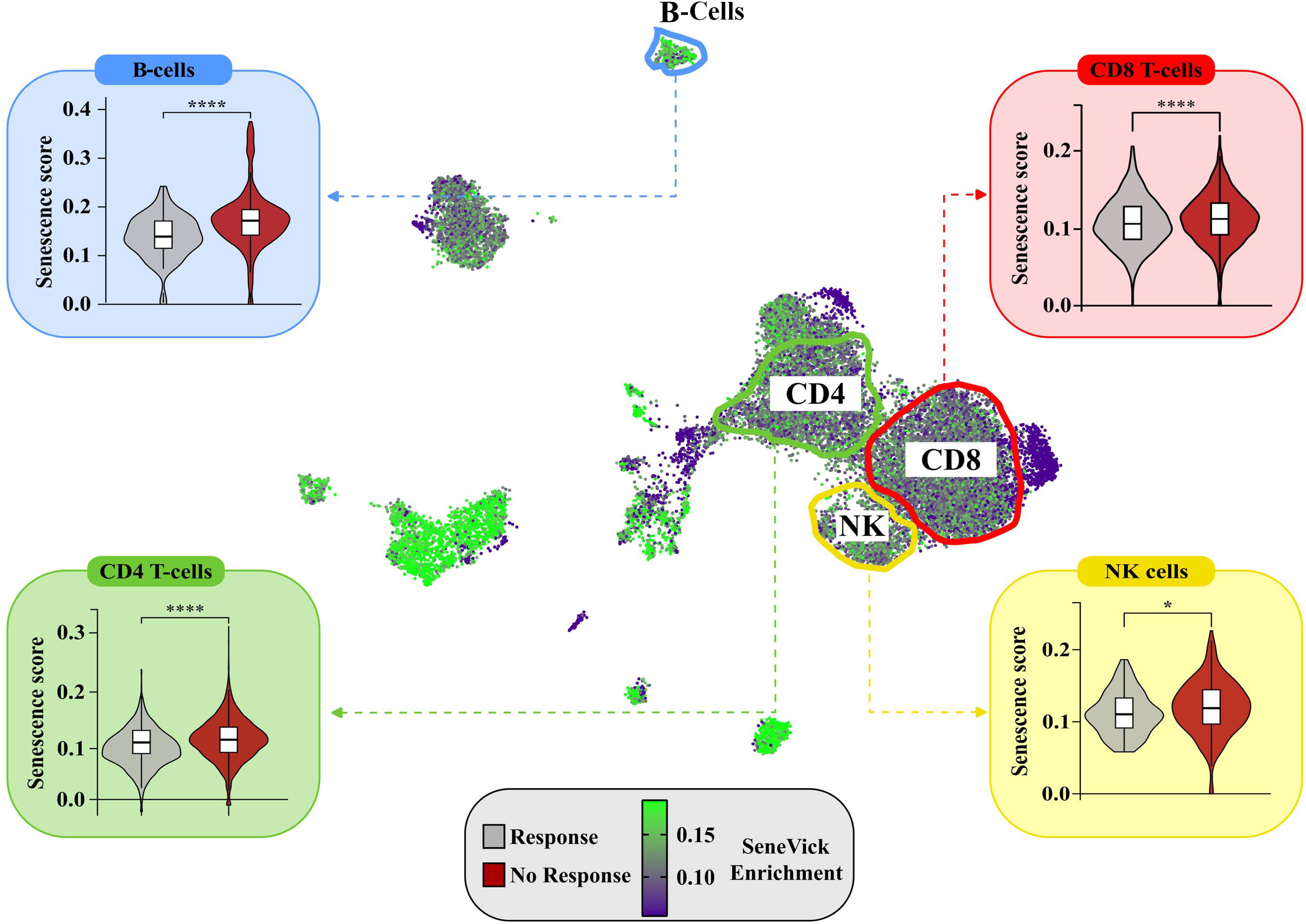
SeneVick implementation in an *in silico* melanoma dataset reveals increased senescent immune cell populations in NRs compared to Rs to immunotherapy. UMAP plot displaying the distribution of the cells from the single-cell RNA sequencing data of the 48 Rs and NRs melanoma patients following immunotherapy, from the GSE115978 (n=30) and GSE120575 (n=18) datasets. The cells are categorized based on SeneVick enrichment (middle panel). Violin plots show significant SeneVick enrichment in CD4+ and CD8+ T-cells, NK and plasma cells of NRs versus Rs (peripheral panels). To visualize the senescence scores, the scores of the cells below the 10th percentile and the ones above the 90th percentile were clipped, in order to reduce the influence of the outliers on the color scale. Nominal *p* values were calculated via Wilcoxon Test.: CD8+ T-cells (p<6.4e-08), CD4+ T-cells (p<3.4e-06), Plasma cells (p<3.6e-08) and NK cells (p<0.016). \**P*1<10.05, \*\**P*1<10.01, \*\*\**P*1<10.001, \*\*\*\**P*1<10.0001

The second approach included analysis of *ex vivo* clinical material from melanoma patients that received single or combined immunotherapy, as monotherapy (**Table S3**). Particularly, peripheral blood cells (PBMCs) from Rs and NRs patients were obtained and senescence was assessed via flow cytometry, using GLF16, an essential component of the SDA^3–5^. This approach was favored as circulating immune cells have been demonstrated to reflect to a large extend the tumor infiltrating ones and provide a reliable snapshot of the TME^51^. As expected, PBMCs originating from NRs patients exhibited significantly higher senescence particularly in the CD4+ and CD8+ T-, and B-cell subtypes compared to Rs (**Figure 5, Figures S5-S6**). This was also the case for NK cells, though the lack of statistical significance was probably due to the low number of *ex vivo* samples. Increased senescence in CD4+ and CD8+ T-, and B-cells of NRs was subsequently confirmed in corresponding tissue biopsies from these patients using the SDA, additionally suggesting that NRs exert considerably increased immune cell senescence, in relation to Rs (**Figure 5, Figure S7**). No association of immune senescence in NRs with clinical features presented in **Table S3** was identified. Overall, the observations from the *in silico* and experimental analysis robustly support that NRs to immunotherapy can be distinguished from Rs based on their immune cell senescence status, irrespective of their age.

**Figure 5:**
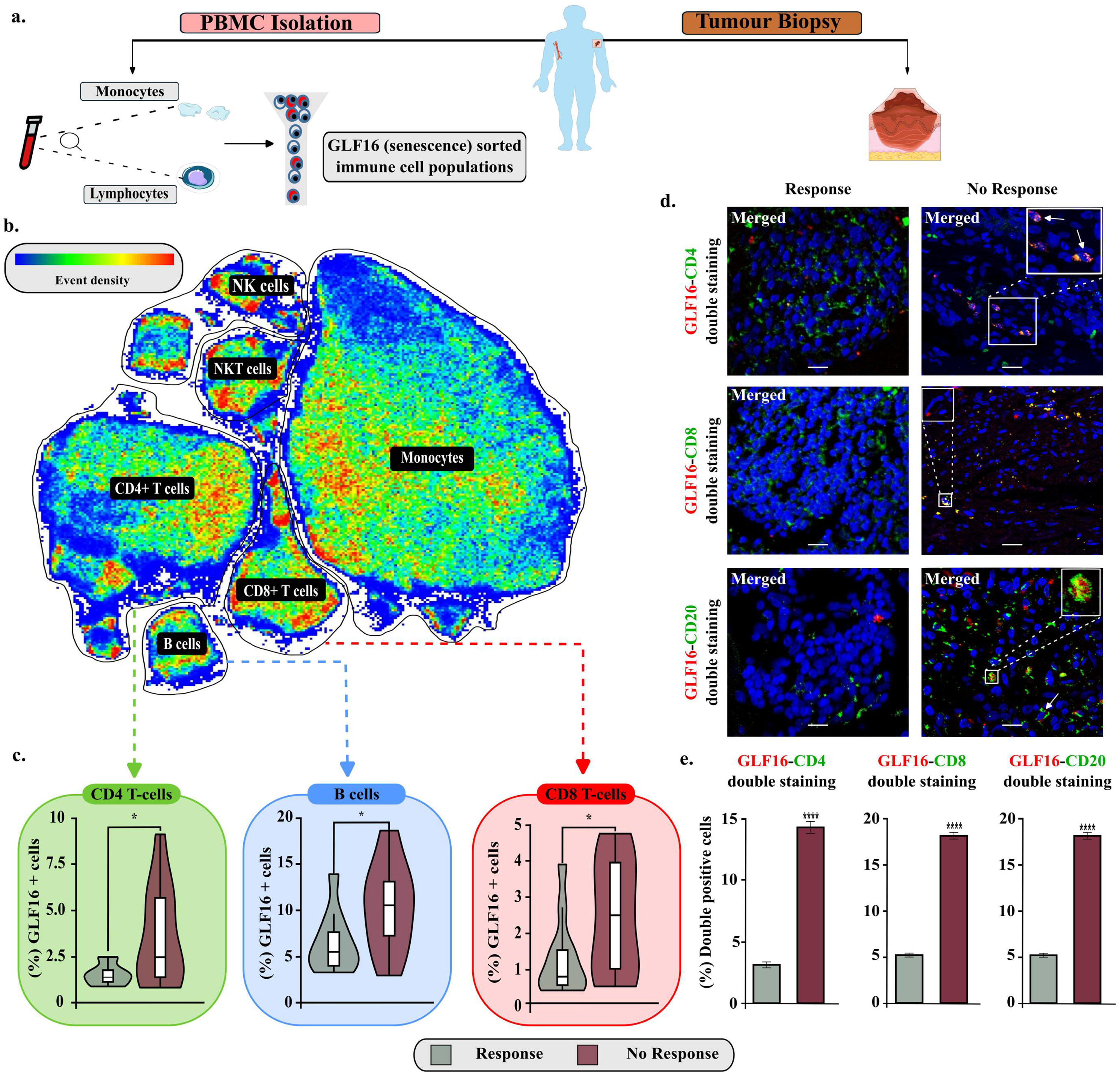
Senescence assessment in an *ex vivo* melanoma cohort, consisting of peripheral blood and tissues, demonstrates increased immune cell senescence in NRs compared to Rs to immunotherapy. **a.** Schematic overview of the experimental procedure followed to collect peripheral blood samples and tumor lesions from melanoma patients. Blood samples were processed with Ficoll-Paque density gradient centrifugation to isolate peripheral blood mononuclear cells (PBMCs). PBMCs were stained with a panel of fluorochrome-conjugated antibodies (**Table S4**) GLF16, and further analyzed with flow cytometry to assess GLF16+ (senescent) immune cell populations. In turn, tissue samples were double stained with GLF16 and immune cell markers to assess immune cell senescence. **b.** Representative t-distributed stochastic neighbor embedding (t-SNE) plot showing the clusters of the major PBMC subsets, i.e., CD4+ T cells, CD8+ T cells, B cells, NK cells, NKT cells, and monocytes. Color coding reflects event density (blue, low; red, high). **c.** Violin plots display the percentages of GLF16+ (senescent) CD4+ and CD8+ T-cells and B-cells, of responders and non-responders, \**P*< 0.05. **d.** Representative images of double staining of cells with the senescence marker GLF16 (red) and CD4 (green), CD8 (green) and CD20 (green). DAPI counterstain. Scale bar: 30μm. The images were quantified using ImageJ. **e.** Quantification of images presented in d. Statistical analysis was performed employing unpaired t-test. The data obtained represent means ± standard deviation. *P*1<10.05, \*\**P*1<10.01, \*\*\**P*1<10.001, \*\*\*\**P*1<10.0001. Objective 20×, Scale bar: 30μm. Melanoma biopsy, human and vessel icons were provided by Servier Medical Art (https://smart.servier.com/), licensed under CC BY 4.0 (https://creativecommons.org/licenses/by/4.0/).”

Moreover, to exclude the possibility that immune checkpoint inhibition could be responsible for senescence observed in the immune cell populations, we implemented SeneVick exclusively in single cell data from melanomas before their treatment. We confirmed that NRs exerted a significantly increased senescence score in CD4+ T-cells (p=6.1*10^-5^), CD8+ T-cells (p=6.5*10^-5^) and NK cells (p=0.011) compared to Rs, prior immunotherapy (**Figure 6**). In plasma cells the difference was not statistically significant, most probably due to the small number of cells of these populations when only the pre-treatment samples are encountered.

**Figure 6:**
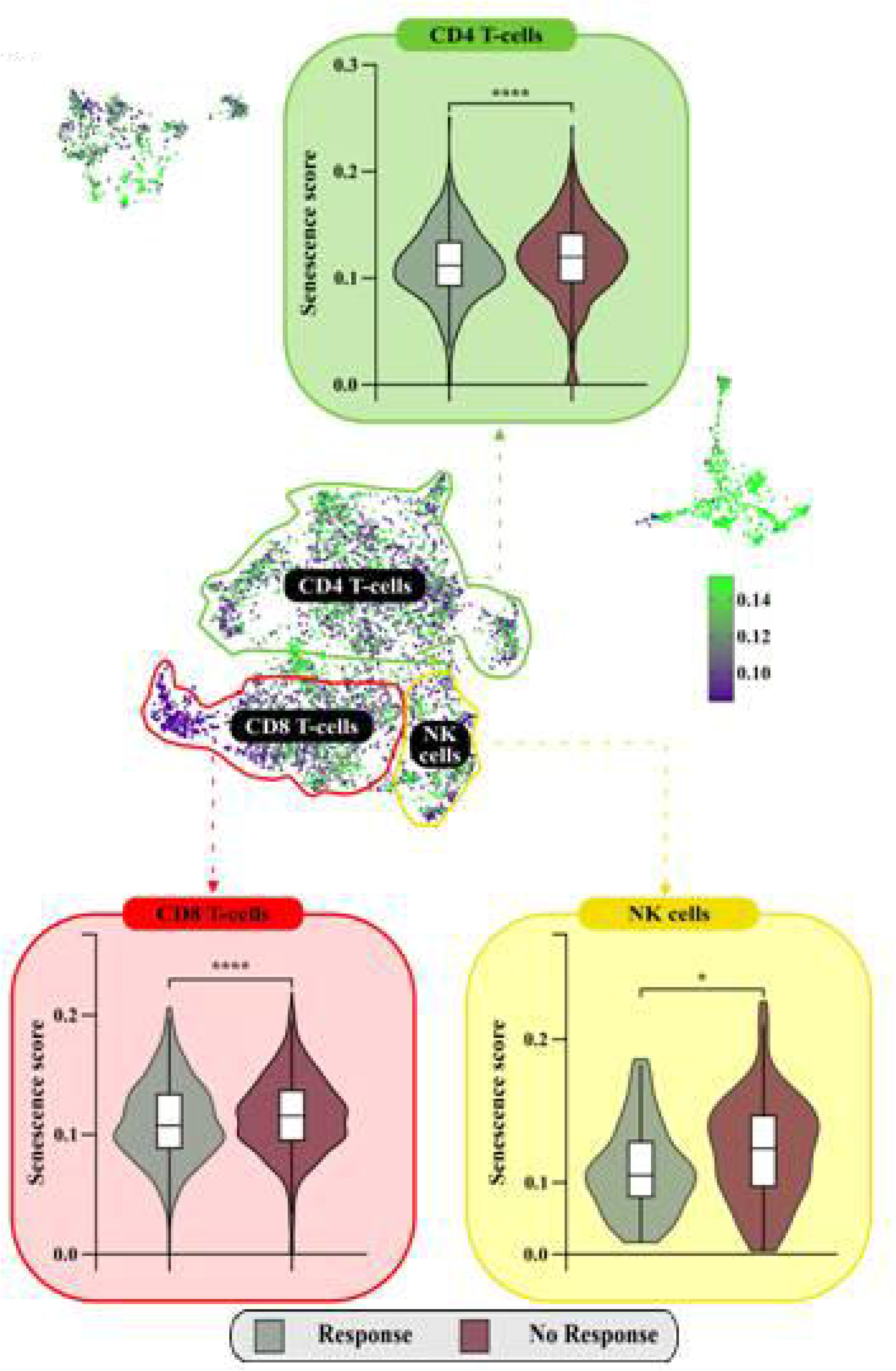
SeneVick identified significantly increased senescence in immune cell populations in NRs vs Rs prior treatment. UMAP plot displaying the distribution of the cells from the single-cell RNA sequencing data of the Rs and NRs melanoma patients prior to immunotherapy treatment, from the GSE115978and GSE120575 datasets. The cells are categorized based on SeneVick enrichment (middle panel). Violin plots show significant SeneVick enrichment in CD4+ and CD8+ T-cells, and NK cells of NRs versus Rs (peripheral panels). To visualize the senescence scores, the scores of the cells below the 20th percentile and the ones above the 80th percentile were clipped, in order to reduce the influence of the outliers on the color scale. Nominal p values were calculated via Wilcoxon Test.: CD8+ T-cells (p=0.00065), CD4+ T-cells (p=6.1e-5) and NK cells (p=0,011). *P1<10.05, **P1<10.01, ***P1<10.001, ****P1<10.0001.

Interestingly, within the CD4+ and CD8+ T-cell subsets of NRs, a population of cells that was neither enriched for SeneVick nor proliferating drew our attention. Further, zooming into this observation, we questioned whether these cells could be exhausted or anergic, as this T-cell states have been previously reported as a source of immune cell dysfunction^52^. In order to examine this issue, we extracted two signatures consisting of the most potent markers identified in the context of exhaustion and anergy respectively (**Table S5**) and applied them in the CD4+ and CD8+ T-cell compartment of our melanoma cohort. Indeed, cell populations negative for senescence, anergy and proliferation exerted an increased enrichment of the “exhaustion” signature while those showing SeneVick enrichment were simultaneously negative for exhaustion, anergy and proliferation (**Figure 7**). These observations highlight the fidelity of SeneVick in discriminating cellular senescence from other dysfunctional cell states within the immune cell compartment, allowing thus to uncover its role not only in cancer but also in various other diseases.

**Figure 7:**
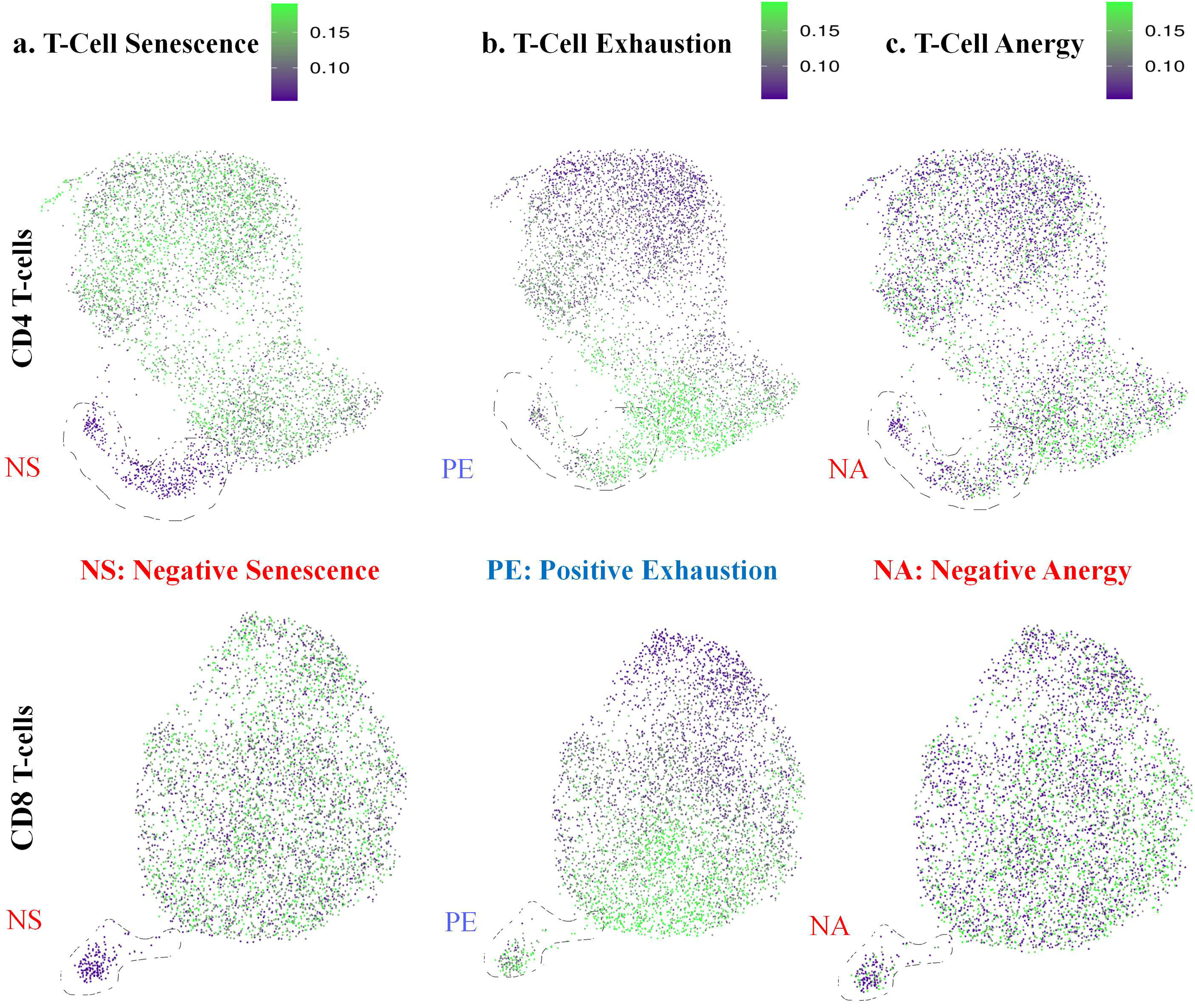
SeneVick effectively discriminates Senescence from Exhaustion and Anergy in the melanoma TME. UMAP plot displaying the distribution of the cells from the single-cell RNA sequencing data of the CD4+ and CD8+ T-cells of the 48 Rs and NRs melanoma patients and categorizing cells after the implementation of SeneVick (**a**), the T-Cell anergy signature (**b**) and the T-Cell exhaustion signature (**c**) As outlined, a subset of non-proliferating CD4+ and CD8+ T-cells in NRs exerts absence of enrichment of SeneVick and the T-cell anergy signature while exhibiting a T-cell exhaustion phenotype. To visualize the signature scores, the scores of the cells below the 10th percentile and the ones above the 90th percentile were clipped, in order to reduce the influence of the outliers on the color scale.

Lastly, in order to gain further insights into the alterations that senescence imposes in the TME of NRs, we analyzed the intercellular interplay of the immune cell subpopulations in Rs versus NRs. Particularly, in the formerly analyzed melanoma scRNA dataset of Rs and NRs patients we applied a cell communication analysis using CellChat^44^. The results showed numerous differences of lig- and-receptor communication probabilities among the immune cell types previously encountered (CD8+ and CD4+ T-cells, NK cells, and plasma cells, **Figure 8a**). These findings suggest that cellular senescence is linked with deregulated immune cell-cell communication in the TME of NRs, leading to dysfunctional immune responses and treatment failure (**Figure 8b**).

**Figure 8:**
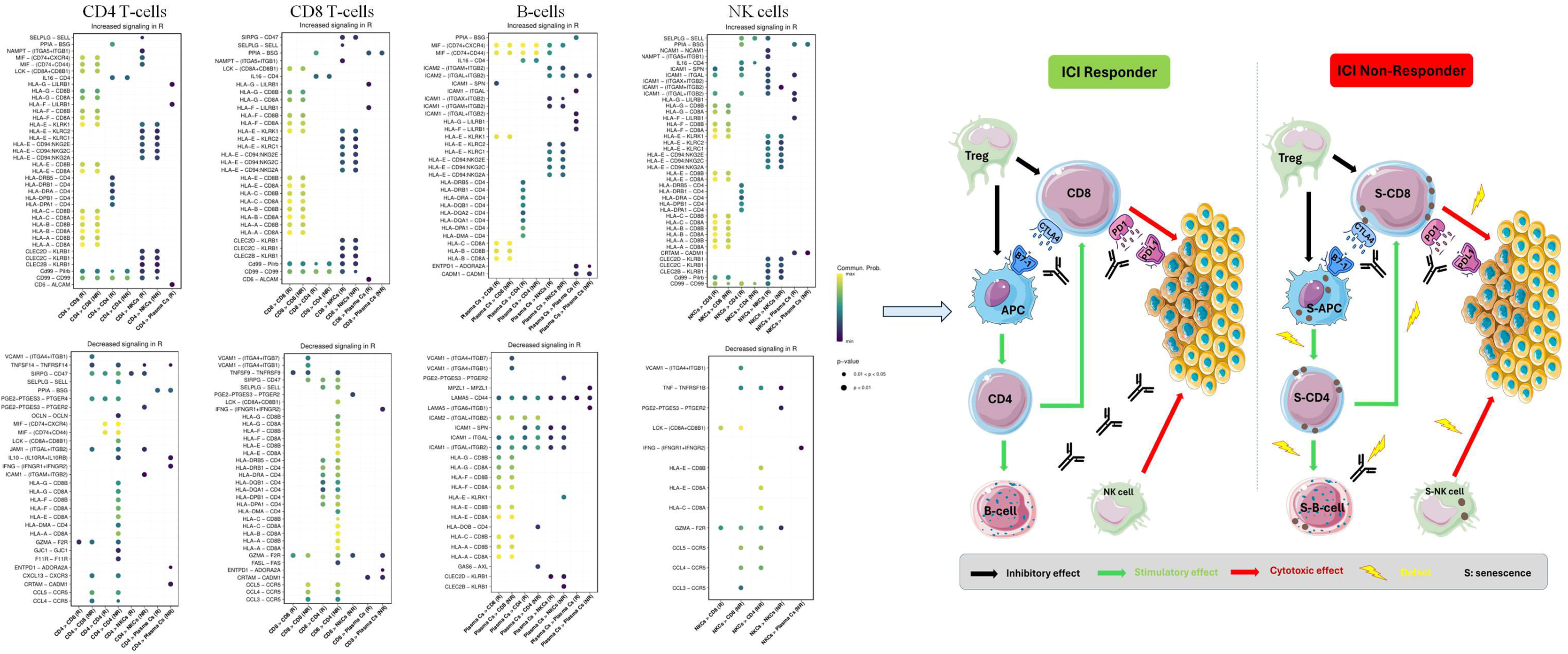
Analysis of the intercellular interplay of the immune cell subpopulations in Rs versus NRs melanoma patients following immunotherapy. **a)** Each panel displays the Ligand-Receptor (LR) interactions with differential strengths between R-NR for different ligand-sender cell types (A: CD4+ T-cells, B: CD8+ T-cells, C: plasma cells, D: NK cells). Each cell type exhibits increased (top) and decreased (bottom) interactions with different immune cell types in Rs patients, compared to non-responding ones. Rows indicate the specific Ligand-Receptor pairs, while columns represent sender–receiver cell type pairs, along with the response status (R or NR) in which the interaction was detected. The color of each dot indicates the interaction strength and the size of the statistical significance. **b)** Schematic illustration representing the altered immune cell interactions in NRs compared to Rs following immunotherapy. Cell icons were provided by Servier Medical Art (https://smart.servier.com/), licensed under CC BY 4.0 (https://creativecommons.org/licenses/by/4.0/).”

## 4. Discussion

Immunotherapy has undoubtedly provided major benefits towards cancer treatment in the last decades, though in a significant portion of patients the treatment yields limited or even no responses^16^. Cancer-related immunodeficiency has arisen as an important determinant of this outcome^53–55^. The latter has been linked to the accumulation of dysfunctional immune cell populations within the TME, such as exhausted and anergic ones while the contribution of cellular senescence is still poorly understood^56–59^. In the context of immunity, senescence has often been inaccurately assessed due to application of debatable and non specific markers^5^. As a consequence, the term immune cell senescence has been misused, its contribution in shaping the TME is still vague and probably underestimated, and its association with the clinical course remains largely unknown^5^. In line with this notion, given that the senescence phenotype is complex and highly heterogeneous, the development of new approaches to assess it more accurately complementing existing or evolving tools, will enhance our understanding on the cellular and molecular mechanisms involved, and its impact in human diseases and clinical outcomes^13,60^.

The current study investigated whether cellular senescence is a source of immune cell dysfunction and is involved in the response to immunotherapy by applying two approaches that complement each other, an *in silico* one as well as a guideline senescence detecting pipeline (SDA), in melanoma patients. Melanoma was selected, as this type of malignancy is commonly treated with immunotherapy as a first option, especially in advanced stages. Moreover, this type of treatment, as shown (**Figure 6**) is not linked to therapy-induced senescence in the immune cell compartment, in contrast to what is observed in traditional ones (irradiation or chemotherapy) that cause DNA damage induced senescence and thus did not influence our observations^61,62^. Regarding the first approach, we initially extensively validated SeneVick, a senescence molecular signature comprising 100 genes that we recently extracted, allowing the discrimination of senescence from aging in the liver^14^. The signature turned out highly sensitive and specific in discriminating non-senescent cells from senescent ones, irrespective of senescence type, tissue origin or species, even upon low senescence levels (**Figures 2-3, Figures S2-S3**). Of note, SeneVick is not a stochastic gene set, but rather a highly structured and evolutionarily curated network of genes likely acting in coordination to mediate key aspects of cellular senescence. The combination of rare protein domains, motif periodicity, and chromosomal distribution patterns not only provides insights into the mechanistic underpinnings of senescence but also hints at the potential translational value of SeneVick in biomarker development, aging research, and therapeutic interventions targeting age-related pathologies.

We next exploited this specific and potent senescence detecting tool, as the first approach, to address the main query of our investigation related to the potential involvement of tumor-related immune cell senescence in the outcome of immunotherapy. By implementing SeneVick retrospectively in a sc-RNA dataset from 48 melanomas treated exclusively with immune checkpoint inhibitors (PD1, CTLA4 or combined inhibition) we observed a considerable signature enrichment in the immune cell populations of NRs, comprising of CD4+ and CD8+ T-cells, NK and plasma cells, that was independent of patient’s age (**Figure 4**). Subsequently, as the second approach, we verified the presence of senescent immune populations in peripheral blood cells (PBMCs) from 11 Rs and 13 NRs melanoma patients following the SDA via flow cytometry^3–5^ (**Table S4**). Indeed, PBMCs originating from NRs patients, particularly CD8+ and CD4+ T-cells, and B-cells exhibited remarkably higher senescence compared to those from Rs (**Figure 5**). This was also the case for NK cells, though the differences were not statistically significant. Interestingly, higher senescence levels were confirmed in these immune cell types within the corresponding tumour lesions of the NR patients compared to Rs (**Figure 5**). These observations were also in line with those that emerged following SeneVick implementation in the *in silico* dataset. Of note, although senescent populations identified in the peripheral blood were quantitatively lower, they reflected qualitatively those in the TME, confirming the value of circulating immune cells as a reliable setting to assess cellular senescence in patients^63,64^ (**Figure 5, Figure S7**).

The above findings underscore the importance of immune cell senescence in driving responsiveness to immunotherapy in melanoma and that the tools implemented herein in a complementary manner are highly efficient to identify those patients who are likely not to respond according to their senescence status. While indications in experimental models support that tumor cell senescence might influence the outcome of immunotherapy our study unveils for the first time the role of immune cell senescence within the TME in responsiveness to such an intervention in melanoma^65^. Interestingly, the *in silico* analysis identified a subset of non-proliferating CD4+ and CD8+ T-cells among NRs where SeneVick signature was not enriched (**Figure 7**). These cells were found to exhibit a T-cell exhaustion signature and were devoid of anergy markers. In line with this notion, NRs patients of the clinical melanoma cohort with the lowest senescence (GLF16) indices exerted characteristically the highest exhaustion levels. These findings denote the reliability of SeneVick and GLF16 staining in discerning cellular senescence from other dysfunctional T-cell states within the TME, a task that so far has been really challenging in phenotypic analyses due to significant overlapping of the applied markers^5,27^. Moreover, numerous differences of intercellular communication among the immune cell populations (CD-8+ and CD4+ T-cells, NK cells and plasma cells) between Rs and NRs, were identified (**Figure 8a**). The latter implies that cellular senescence is linked with deregulated immune cell to cell communication in the TME of NRs in favor of dysfunctional immune responses (**Figure 8b**). Collectively, our observations support a cell senescence-mediated immune suppressive environment, imposing resistance to immunotherapy (**Figure 8b**). The latter is characterized by senescence in CD8+ T-cells, and NK cells leading to loss of their cytotoxic activity, while senescence in CD4+ cells that exert a multifaceted role in cancer immunity, leads to a diminished pool of functional T-cells incapable of responding to new antigens^66–68^. Senescent cell populations, particularly CD8+ and CD4+ T-cells, can secrete a variety of SASP factors (pro-inflammatory cytokines, chemokines and growth factors), that can remodel the immune landscape and influence the TME towards tumor progression^69–71^. In this context, tumor infiltrating senescent T lymphocytes can affect B-cell activation and their subsequent differentiation into plasma cells, ultimately leading to a deficiency in antibody production and inefficient adaptive responses ^69–71^. The latter is also a potential outcome of B-cell senescence identified in our analysis in NRs.

While there seems to be a relation between aging and immune senescence^71^, our findings support that cancer cells can shape a microenvironment to promote immune cell senescence as a strategy for immune evasion, independent of patient’s age, addressing thus a debatable matter. Potential mechanisms involved are summarized in Figure S8. Nevertheless, these mechanisms should be regarded cautiously, as markers applied for senescence identification in these studies can also be evident in other immune cell dysfunctional states^5^.

Given the plasticity of immune cells and that anergy or exhaustion reflect in principle progressive and irreversible states acquired upon chronic infections or cancer, immune cell senescence emerges as an attractive option to rejuvenate the immune system in order to restore its functionality, increasing thus the efficacy of immunotherapy. In fact, strategies for immune cell reinvigoration that target the above molecular mechanisms and pathways as well as elimination of the toxic and immunosuppressive senescent cell compartment in the TME are gaining increased attention^5,72^. Regarding the latter, a recently reported innovative senolytic platform that allows for selective removal of senescent cells without side effects paves the way^73^. This advancement underscores the importance of investigations such as the current one that exploits efficient senescence detecting tools to characterize patients according to their senescence status which drives their responsiveness to therapy.

---

**Figure S1: Zooming into the SeneVick signature** a) Pie chart illustrating the distribution of the 100 genes across major functional categories identified through gene ontology (GO) enrichment and biochemical pathway analysis. Genes were grouped into representative clusters including cell cycle regulation, DNA damage response, chromatin remodeling, inflammatory signaling, metabolic processes, and secretory phenotype regulation. Categories were defined based on overrepresented GO terms and curated pathway annotations from KEGG, Reactome, and GeneCards. The chart highlights the relative abundance of genes involved in each biological process, reflecting the multifaceted molecular landscape of cellular senescence.

b) Venn diagram showing overlapping genes across SeneVick and the two most robust senescence signatures.

**Figure S2: SeneVick efficiently detects senescence across tissues and upon age** Violin Plots demonstrate significant SeneVick enrichment in different cell types (**a**) and upon time (**b**) in aged vs young mice (n=19 male and n=11 female, GSE132042).

Two data sets were compared with the Wilcoxon test, \**P*1<10.05, \*\**P*1<10.01, \*\*\**P*1<10.001, \*\*\*\**P*1<10.0001

**Figure S3: SeneVick exerts increased sensitivity and specificity in demarcating senescent cells from non-senescent ones compared to other senescence detecting signatures a.** Right panel: Indication of the different timepoints on UMAP plot of the scRNA data from human fibroblasts in the GSE226225 dataset.

**b.** UMAP plot categorizing cells based on SeneVick, SenMayo and FridMan enrichment upon time (days) in the GSE226225 dataset showing the scRNA data of human fibroblasts (GSE226225) representing the enrichment of the three signatures upon time following treatment.

**c.** Timepoint analysis (p<0,05) showing the gradual enrichment of the signatures across days following etoposide treatment from the scRNA data of GSE226225 and demonstrating the occurring breakpoints (day 1). The enrichment levels of SeneVick (**top**), SenMayo (**middle**) and FridMan (**bottom**) in human fibroblasts, in which the induction of senescence was accomplished with different senescent inducers (IR and ETO) were compared to proliferative fibroblasts. Significance was assessed by Wilcoxon Test, p < 2.22e-16. \**P*1<10.05, \*\**P*1<10.01, \*\*\**P*1<10.001, \*\*\*\**P*1<10.0001

**Figure S4: Workflow for the integration of scRNA patients’ data from the two melanoma studies** Schematic illustration of data preprocessing (**Box 1**), quality control (**Box 2**), normalization (**Box 3**), and cell annotation for the integration of the scRNA data of the Rs and NRs melanoma patients following immunotherapy from the GSE115978 (n=30) and GSE120575 (n=18) datasets.

**Figure S5: Gating strategy followed for the identification of peripheral blood mononuclear cell (PBMC) subpopulations and the assessment of GLF16+ (senescent) cells.** Exclusion of acquisition artefacts was performed by plotting events over time. Single cells were selected by sequential gating on FSC-A / FSC-H and SSC-A / SSC-H dot plots. Live cells were identified with BD Horizon™ Fixable Viability Stain 570. In a CD14 / CD16 plot, monocytes were gated and further classified into classical (CD14++CD16-), intermediate (CD14++CD16+), and non-classical (CD14+CD16++). CD14-cells were plotted in a CD3 / CD19 plot to identify B-cells, further classified in a CD27 / CD38 plot to plasma cells (CD27+CD38+), naïve B-cells (CD27-CD38-) and memory B-cells (CD27+CD38-). In the same plot, gated CD3-CD19 negative cells were shown in a CD16 / CD56 plot, and gated NK cells were further classified as CD56bright, CD56+ and CD56dim. Finally, T / NKT cells were distinguished in a CD56 / SSC-A plot, and T-cells were further shown in a CD4 / SSC-A plot to gate CD8+ and CD4+ T-cells. Tregs were identified as CD4+CD127-CD25high. CD4+, CD8+ T and NKT cells were subdivided in CD27 / CD28 plots. Percentages of GLF16+ cells in CD4+ and CD8+ T-cells, NK and B-cells are shown as inserts.

**Figure S6: GLF16+ (senescent) cell % percentages of immune subsets of Rs and NRs melanoma patients. a.** Workflow depicting cellular senescence assessment in circulating CD4+ and CD8+ T-cells and B-cells from melanoma patients by applying the senescence detecting algorithm (SDA).

**b.** Representative dot plots from 4 melanoma patients, showing the percentages of GLF16+ cells (in inserts) in CD4+ and CD8+ T-cells, and B-cells isolated from two Rs (Responder 1 and 2) and two NRs (Non-Responder 1 and 2).

Human and vessel icons were provided by Servier Medical Art (https://smart.servier.com/), licensed under CC BY 4.0 (https://creativecommons.org/licenses/by/4.0/).”

**Figure S7: Serial section analysis reveals increased immune cell senescence in the TME of NRs compared to Rs to immunotherapy** Representative images of serial section analysis in melanoma lesions stained with the senescence detecting reagent GL13 and markers for CD4+ and CD8+ T-cells, and B-cells. Rs (upper panel) depict lower senescence in CD4, CD8 and CD20 cells within the TME compared to NRs (lower panel: insets and red arrows) Objective 10× (1^st^ and 3^rd^ line), 40× (2^nd^ and 4^th^). Scale bars: 30 μm and 60 μm, respectively.

**Figure S8: Overview of cancer driven mechanisms inducing immune cell senescence in the TME** Schematic illustration of putative mechanisms involved in cancer promoted immune cell senescence in the tumor microenvironment. Due to cancer cell’s accelerated growth and metabolism the TME is characterized by hypoxia, high levels of reactive oxygen (ROS) and nitrogen (RNS) species and lipids that solely or in concert induce DNA damage response, eventually triggering immune cell senescence^75^. Interestingly, tumor cells have been reported to induce senescence in T-cells by c-AMP delivery or by transferring mitochondria with mtDNA mutations^76,77^. Moreover, tumor-derived immunoglobulin-like transcript 4 (ILT4), an inhibitory molecule of the immunoglobulin superfamily, has been shown to induce T-cell senescence via activation of ERK1/2 MAPK signaling^78^. Tumor-associated Tregs can also induce T-cell senescence in responding naïve/effector T-cells, by promoting mitochondrial disruption and p38/ERK1/2 MAPK signaling activation as well as via increased glucose consumption and metabolic competition^79^.

Cell icons were provided by Servier Medical Art (https://smart.servier.com/), licensed under CC BY 4.0 (https://creativecommons.org/licenses/by/4.0/).”

## Acknowledgements

The authors would like to acknowledge Dr Sarantis Gagos, Dr Aikaterini Polyzou, Dr Alexandros Athanasopoulos and Dr Athanasios Korogiannos for their valuable contribution in preparing the manuscript.

## Ethics declarations

Ethics approval and consent to participate

All procedures performed in this study involving human participants were in accordance with the ethical standards of the…

The study was reviewed and approved by Bioethics Committee…, and informed consent was obtained from the patients.

## Consent for publication

All authors contributed to the writing of the manuscript. All authors approved the final version of the manuscript.

## References

1. Gorgoulis V, Adams PD, Alimonti A, et al. Cellular Senescence: Defining a Path Forward. Cell. 2019;179(4):813–827. doi:10.1016/j.cell.2019.10.005

2. Lawless C, Wang C, Jurk D, Merz A, Zglinicki T von, Passos JF. Quantitative assessment of markers for cell senescence. Exp Gerontol. 2010;45(10):772–778. doi:10.1016/j.exger.2010.01.018

3. J K, B W, Sm B, et al. Algorithmic assessment of cellular senescence in experimental and clinical specimens. Nat Protoc. 2021;16(5). doi:10.1038/s41596-021-00505-5

4. M O, J CA, Pd A, et al. Guidelines for minimal information on cellular senescence experimentation in vivo. Cell. 2024;187(16). doi:10.1016/j.cell.2024.05.059

5. Magkouta S, Markaki E, Evangelou K, Petty R, Verginis P, Gorgoulis V. Decoding T cell senescence in cancer: Is revisiting required? Semin Cancer Biol. 2025;108:33–47. doi:10.1016/j.semcancer.2024.11.003

6. Bartkova J, Rezaei N, Liontos M, et al. Oncogene-induced senescence is part of the tumorigenesis barrier imposed by DNA damage checkpoints. Nature. 2006;444(7119):633-637. doi:10.1038/nature05268

7. Evangelou K, Veroutis D, Paschalaki K, et al. Pulmonary infection by SARS-CoV-2 induces senescence accompanied by an inflammatory phenotype in severe COVID-19: possible implications for viral mutagenesis. Eur Respir J. Published online January 27, 2022. doi:10.1183/13993003.02951-2021

8. Senescent cells in giant cell arteritis display an inflammatory phenotype participating in tissue injury via IL-6-dependent pathways - ScienceDirect. Accessed July 27, 2025. https://www.sciencedirect.com/science/article/abs/pii/S0003496724003625?via%3Dihub

9. Galanos P, Vougas K, Walter D, et al. Chronic p53-independent p21 expression causes genomic instability by deregulating replication licensing. Nat Cell Biol. 2016;18(7):777–789. doi:10.1038/ncb3378

10. Zampetidis CP, Galanos P, Angelopoulou A, et al. A recurrent chromosomal inversion suffices for driving escape from oncogene-induced senescence via subTAD reorganization. Mol Cell. 2021;81(23):4907–4923.e8. doi:10.1016/j.molcel.2021.10.017

11. V M, K E, Pvs V, et al. Senescence and senotherapeutics: a new field in cancer therapy. Pharmacol Ther. 2019;193. doi:10.1016/j.pharmthera.2018.08.006

12. Al C, A F, Ja AP, et al. Towards frailty biomarkers: Candidates from genes and pathways regulated in aging and age-related diseases. Ageing Res Rev. 2018;47. doi:10.1016/j.arr.2018.07.004

13. Y S, Q L, Jl K. Targeting senescent cells for a healthier longevity: the roadmap for an era of global aging. Life Med. 2022;1(2). doi:10.1093/lifemedi/lnac030

14. K E, K B, P P, et al. In Situ and In Silico Methods for Senescence Identification in Human Liver Diseases. Methods Mol Biol Clifton NJ. 2025;2906. doi:10.1007/978-1-0716-4426-3_1

15. C B, A T, E R, et al. Safety profiles of anti-CTLA-4 and anti-PD-1 antibodies alone and in combination. Nat Rev Clin Oncol. 2016;13(8). doi:10.1038/nrclinonc.2016.58

16. Ad W, Jm F, Mj L. A guide to cancer immunotherapy: from T cell basic science to clinical practice. Nat Rev Immunol. 2020;20(11). doi:10.1038/s41577-020-0306-5

17. M P, A S. CD8+ T cell differentiation and dysfunction in cancer. Nat Rev Immunol. 2022;22(4). doi:10.1038/s41577-021-00574-3

18. N W, M R, C A, et al. Single-cell transcriptomic analysis uncovers diverse and dynamic senescent cell populations. Aging. 2023;15(8). doi:10.18632/aging.204666

19. A single-cell transcriptomic atlas characterizes ageing tissues in the mouse. Nature. 2020;583(7817). doi:10.1038/s41586-020-2496-1

20. S Y, Se C, Y K, et al. Decontamination of ambient RNA in single-cell RNA-seq with DecontX. Genome Biol. 2020;21(1). doi:10.1186/s13059-020-1950-6

21. Saul D, Kosinsky RL, Atkinson EJ, et al. A new gene set identifies senescent cells and predicts senescence-associated pathways across tissues. Nat Commun. 2022;13(1):4827. doi:10.1038/s41467-022-32552-1

22. Al F, Ma T. Critical pathways in cellular senescence and immortalization revealed by gene expression profiling. Oncogene. 2008;27(46). doi:10.1038/onc.2008.213

23. Kolberg L, Raudvere U, Kuzmin I, Adler P, Vilo J, Peterson H. g:Profiler—interoperable web service for functional enrichment analysis and gene identifier mapping (2023 update). Nucleic Acids Res. 2023;51(W1):W207–W212. doi:10.1093/nar/gkad347

24. Bt S, M H, J Q, et al. DAVID: a web server for functional enrichment analysis and functional annotation of gene lists (2021 update). Nucleic Acids Res. 2022;50(W1). doi:10.1093/nar/gkac194

25. Ikotun AM, Ezugwu AE, Abualigah L, Abuhaija B, Heming J. K-means clustering algorithms: A comprehensive review, variants analysis, and advances in the era of big data. Inf Sci. 2023;622:178–210. doi:10.1016/j.ins.2022.11.139

26. Georgakopoulou E, Tsimaratou K, Evangelou K, et al. Specific lipofuscin staining as a novel biomarker to detect replicative and stress-induced senescence. A method applicable in cryo-preserved and archival tissues. Aging. 2012;5(1):37–50. doi:10.18632/aging.100527

27. S M, D V, A P, et al. A fluorophore-conjugated reagent enabling rapid detection, isolation and live tracking of senescent cells. Mol Cell. 2023;83(19). doi:10.1016/j.molcel.2023.09.006

28. Magkouta S, Veroutis D, Pousias A, et al. One-step rapid tracking and isolation of senescent cells in cellular systems, tissues, or animal models via GLF16. STAR Protoc. 2024;5(1):102929. doi:10.1016/j.xpro.2024.102929

29. Pulsed Electromagnetic Fields (PEMFs) Trigger Cell Death and Senescence in Cancer Cells. Accessed July 27, 2025. https://www.mdpi.com/1422-0067/25/5/2473

30. Fm R, Sk S, V K, M C, Td H, S G. Alternative lengthening of human telomeres is a conservative DNA replication process with features of break-induced replication. EMBO Rep. 2016;17(12). doi:10.15252/embr.201643169

31. A D, Ca D, F S, et al. STAR: ultrafast universal RNA-seq aligner. Bioinforma Oxf Engl. 2013;29(1). doi:10.1093/bioinformatics/bts635

32. Li H, Handsaker B, Wysoker A, et al. The Sequence Alignment/Map format and SAMtools. Bioinformatics. 2009;25(16):2078–2079. doi:10.1093/bioinformatics/btp352

33. D R, J N, Tp S, S D. Normalization of RNA-seq data using factor analysis of control genes or samples. Nat Biotechnol. 2014;32(9). doi:10.1038/nbt.2931

34. Mi L, W H, S A. Moderated estimation of fold change and dispersion for RNA-seq data with DESeq2. Genome Biol. 2014;15(12). doi:10.1186/s13059-014-0550-8

35. G D, Bt S, Da H, et al. DAVID: Database for Annotation, Visualization, and Integrated Discovery. Genome Biol. 2003;4(5). Accessed July 27, 2025. https://pubmed.ncbi.nlm.nih.gov/12734009/

36. Chang W, Cheng J, Allaire JJ, et al. shiny: Web Application Framework for R. Published online July 3, 2025. Accessed July 27, 2025. https://cran.r-project.org/web/packages/shiny/index.html

37. A S, P T, Vk M, et al. Gene set enrichment analysis: a knowledge-based approach for interpreting genome-wide expression profiles. Proc Natl Acad Sci U S A. 2005;102(43). doi:10.1073/pnas.0506580102

38. Sade-Feldman M, Yizhak K, Bjorgaard SL, et al. Defining T cell states associated with response to checkpoint immunotherapy in melanoma. Cell. 2018;175(4):998. doi:10.1016/j.cell.2018.10.038

39. Jerby-Arnon L, Shah P, Cuoco MS, et al. A Cancer Cell Program Promotes T Cell Exclusion and Resistance to Checkpoint Blockade. Cell. 2018;175(4):984. doi:10.1016/j.cell.2018.09.006

40. Aran D, Looney AP, Liu L, et al. Reference-based analysis of lung single-cell sequencing reveals a transitional profibrotic macrophage. Nat Immunol. 2019;20(2):163–172. doi:10.1038/s41590-018-0276-y

41. C DC, C X, Lb J, et al. Cross-tissue immune cell analysis reveals tissue-specific features in humans. Science. 2022;376(6594). doi:10.1126/science.abl5197

42. New response evaluation criteria in solid tumours: revised RECIST guideline (version 1.1) - PubMed. Accessed July 30, 2025. https://pubmed.ncbi.nlm.nih.gov/19097774/

43. M A, Sj C. UCell: Robust and scalable single-cell gene signature scoring. Comput Struct Biotechnol J. 2021;19. doi:10.1016/j.csbj.2021.06.043

44. Jin S, Guerrero-Juarez CF, Zhang L, et al. Inference and analysis of cell-cell communication using CellChat. Nat Commun. 2021;12(1):1088. doi:10.1038/s41467-021-21246-9

45. Schroeder HT, Muller CHDL, Heck TG, Krause M, Paulo Ivo Homem de Bittencourt J. Resolution of inflammation in chronic disease via restoration of the heat shock response (HSR). Cell Stress Chaperones. 2024;29(1):66. doi:10.1016/j.cstres.2024.01.005

46. S S, Bc R, N H, T A, K G, R K. A novel ankyrin repeat-containing gene (Kank) located at 9p24 is a growth suppressor of renal cell carcinoma. J Biol Chem. 2002;277(39). doi:10.1074/jbc.M204244200

47. L K, M F, H H. DNA motifs and sequence periodicities. In Silico Biol. 2006;6(1-2). Accessed July 27, 2025. https://pubmed.ncbi.nlm.nih.gov/16789915/

48. (PDF) Patau Syndrome: Genetic and Epigenetic Aspects. ResearchGate. doi:10.2991/978-94-6463-062-6_32

49. Hs M, A M, V D, et al. Down-syndrome-induced senescence disrupts the nuclear architecture of neural progenitors. Cell Stem Cell. 2022;29(1). doi:10.1016/j.stem.2021.12.002

50. Farmakis SG, Barnes AM, Carey JC, Braddock SR. Solid tumor screening recommendations in trisomy 18. Am J Med Genet A. 2019;179(3):455–466. doi:10.1002/ajmg.a.61029

51. J H, E H, Y W, et al. Circulating immune biomarkers correlating with response in patients with metastatic renal cell carcinoma on immunotherapy. JCI Insight. 2025;10(4). doi:10.1172/jci.insight.185963

52. En K, A S, J D, et al. Dysfunctional states of unconventional T-cell subsets in cancer. J Leukoc Biol. 2024;115(1). doi:10.1093/jleuko/qiad129

53. Wu B, Zhang B, Li B, Wu H, Jiang M. Cold and hot tumors: from molecular mechanisms to targeted therapy. Signal Transduct Target Ther. 2024;9(1):274. doi:10.1038/s41392-024-01979-x

54. Yt L, Zj S. Turning cold tumors into hot tumors by improving T-cell infiltration. Theranostics. 2021;11(11). doi:10.7150/thno.58390

55. Bilotta MT, Antignani A, Fitzgerald DJ. Managing the TME to improve the efficacy of cancer therapy. Front Immunol. 2022;13. doi:10.3389/fimmu.2022.954992

56. Zhang Z, Liu S, Zhang B, Qiao L, Zhang Y. T Cell Dysfunction and Exhaustion in Cancer. Front Cell Dev Biol. 2020;8:17. doi:10.3389/fcell.2020.00017

57. Braun DA, Street K, Burke KP, et al. Progressive immune dysfunction with advancing disease stage in renal cell carcinoma. Cancer Cell. 2021;39(5):632. doi:10.1016/j.ccell.2021.02.013

58. Peralta RM, Xie B, Lontos K, et al. Dysfunction of exhausted T cells is enforced by MCT11-mediated lactate metabolism. Nat Immunol. 2024;25(12):2297–2307. doi:10.1038/s41590-024-01999-3

59. Lee YH, Tsai KW, Lu KC, Shih LJ, Hu WC. Cancer as a Dysfunctional Immune Disorder: Pro-Tumor TH1-like Immune Response and Anti-Tumor THαβ Immune Response Based on the Complete Updated Framework of Host Immunological Pathways. Biomedicines. 2022;10(10):2497. doi:10.3390/biomedicines10102497

60. Tc B, Je V, Ao S, Lr F, E FC. Subcellular structure, heterogeneity, and plasticity of senescent cells. Aging Cell. 2024;23(4). doi:10.1111/acel.14154

61. Mahalingam P, Newsom-Davis T. Cancer immunotherapy and the management of side effects. Clin Med. 2024;23(1):56. doi:10.7861/clinmed.2022-0589

62. Ja E, Ja D, G W, Df J. Therapy-induced senescence in cancer. J Natl Cancer Inst. 2010;102(20). doi:10.1093/jnci/djq364

63. J S, H W, X L, et al. Deep Sequencing of T-Cell Receptors for Monitoring Peripheral CD8+ T Cells in Chinese Advanced Non-Small-Cell Lung Cancer Patients Treated With the Anti-PD-L1 Antibody. Front Mol Biosci. 2021;8. doi:10.3389/fmolb.2021.679130

64. Ma AM, K M, E E. Correlations between Circulating and Tumor-Infiltrating CD4+ T Cell Subsets with Immune Checkpoints in Colorectal Cancer. Vaccines. 2022;10(4). doi:10.3390/vaccines10040538

65. Maggiorani D, Le O, Lisi V, et al. Senescence drives immunotherapy resistance by inducing an immunosuppressive tumor microenvironment. Nat Commun. 2024;15(1):2435. doi:10.1038/s41467-024-46769-9

66. Ys M, Vp T, Da B, Sa R. Reliable Hallmarks and Biomarkers of Senescent Lymphocytes. Int J Mol Sci. 2023;24(21). doi:10.3390/ijms242115653

67. D Z, M B, Ak S. Hallmarks and detection techniques of cellular senescence and cellular ageing in immune cells. Aging Cell. 2021;20(2). doi:10.1111/acel.13316

68. J Y, M L, D H, M Z, X Z. The Paradoxical Role of Cellular Senescence in Cancer. Front Cell Dev Biol. 2021;9. doi:10.3389/fcell.2021.722205

69. Y G, Y L, X L, et al. CD4+ T-Cell Senescence in Neurodegenerative Disease: Pathogenesis and Potential Therapeutic Targets. Cells. 2024;13(9). doi:10.3390/cells13090749

70. Liu Z, Liang Q, Ren Y, et al. Immunosenescence: molecular mechanisms and diseases. Signal Transduct Target Ther. 2023;8(1):200. doi:10.1038/s41392-023-01451-2

71. Wang Y, Dong C, Han Y, Gu Z, Sun C. Immunosenescence, aging and successful aging. Front Immunol. 2022;13. doi:10.3389/fimmu.2022.942796

72. K E, Vg G. Rejuvenating the immune system. Mol Oncol. 2025;19(3). doi:10.1002/1878-0261.13802

73. Magkouta S, Veroutis D, Papaspyropoulos A, et al. Generation of a selective senolytic platform using a micelle-encapsulated Sudan Black B conjugated analog. Nat Aging. 2025;5(1):162–175. doi:10.1038/s43587-024-00747-4

74. Z W, M G, M S. RNA-Seq: a revolutionary tool for transcriptomics. Nat Rev Genet. 2009;10(1). doi:10.1038/nrg2484

75. Gorgoulis VG, Pefani DE, Pateras IS, Trougakos IP. Integrating the DNA damage and protein stress responses during cancer development and treatment. J Pathol. 2018;246(1):12–40. doi:10.1002/path.5097

76. Ye J, Ma C, Hsueh EC, et al. TLR8 signaling enhances tumor immunity by preventing tumorlinduced Tlcell senescence. EMBO Mol Med. 2014;6(10):1294–1311. doi:10.15252/emmm.201403918

77. Ikeda H, Kawase K, Nishi T, et al. Immune evasion through mitochondrial transfer in the tumour microenvironment. Nature. 2025;638(8049):225-236. doi:10.1038/s41586-024-08439-0

78. Wang Z, He L, Li W, et al. GDF15 induces immunosuppression via CD48 on regulatory T cells in hepatocellular carcinoma. J Immunother Cancer. 2021;9(9):e002787. doi:10.1136/jitc-2021-002787

79. Liu X, Hoft DF, Peng G. Senescent T cells within suppressive tumor microenvironments: emerging target for tumor immunotherapy. J Clin Invest. 130(3):1073–1083. doi:10.1172/JCI133679

