## Supplementary figures and images for "Immune cell senescence drives responsiveness to immunotherapy in melanoma"

### Suppl Fig 1

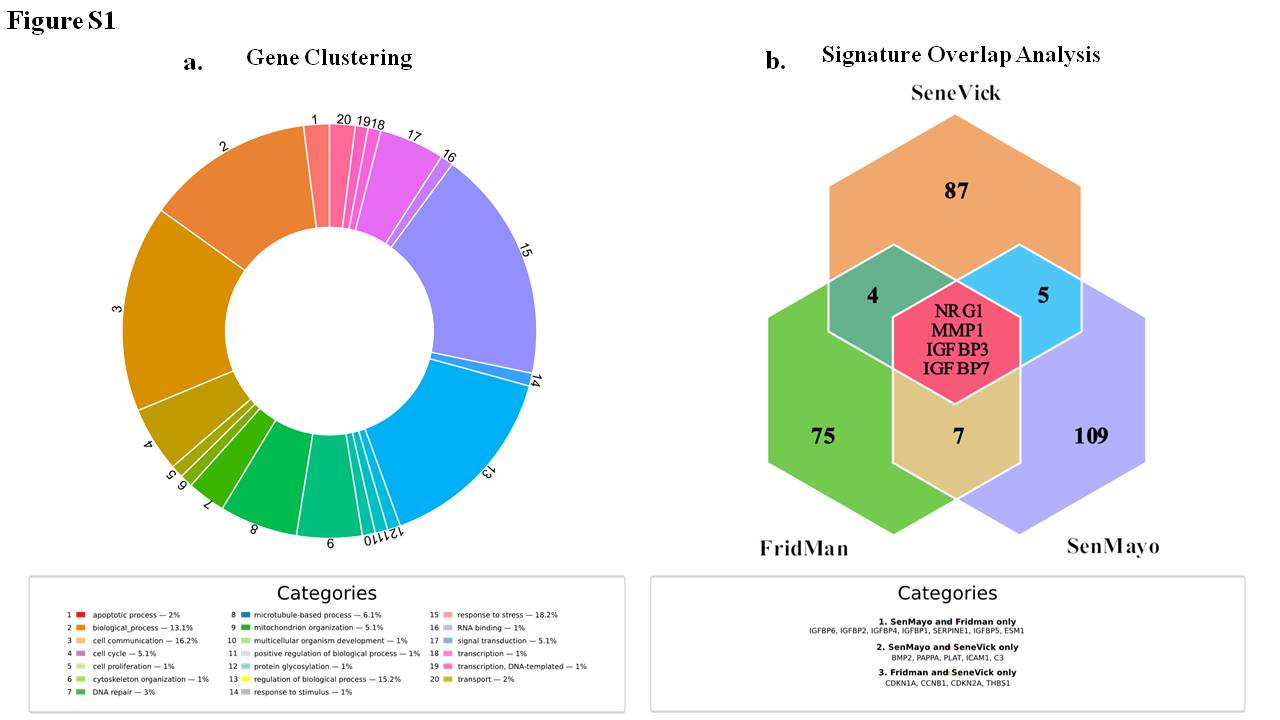

### Suppl Fig 2

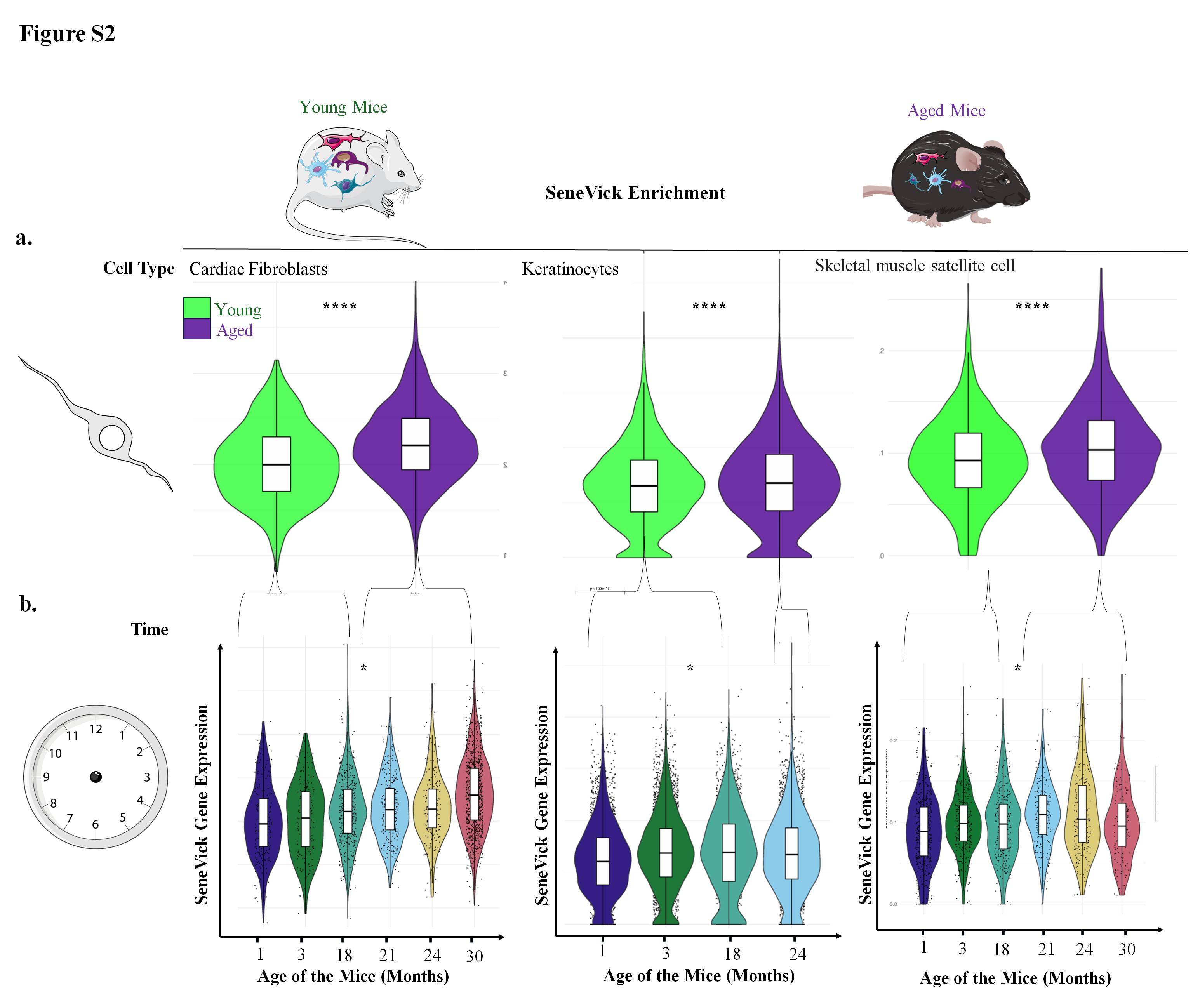

### Suppl Fig 3

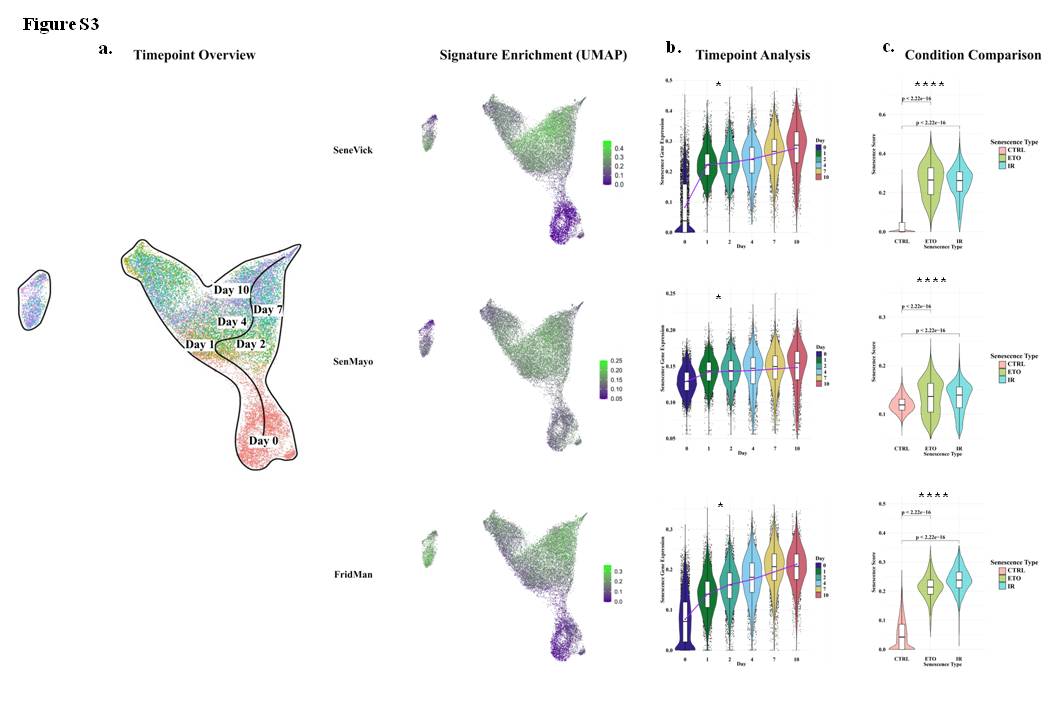

### Suppl Fig 4

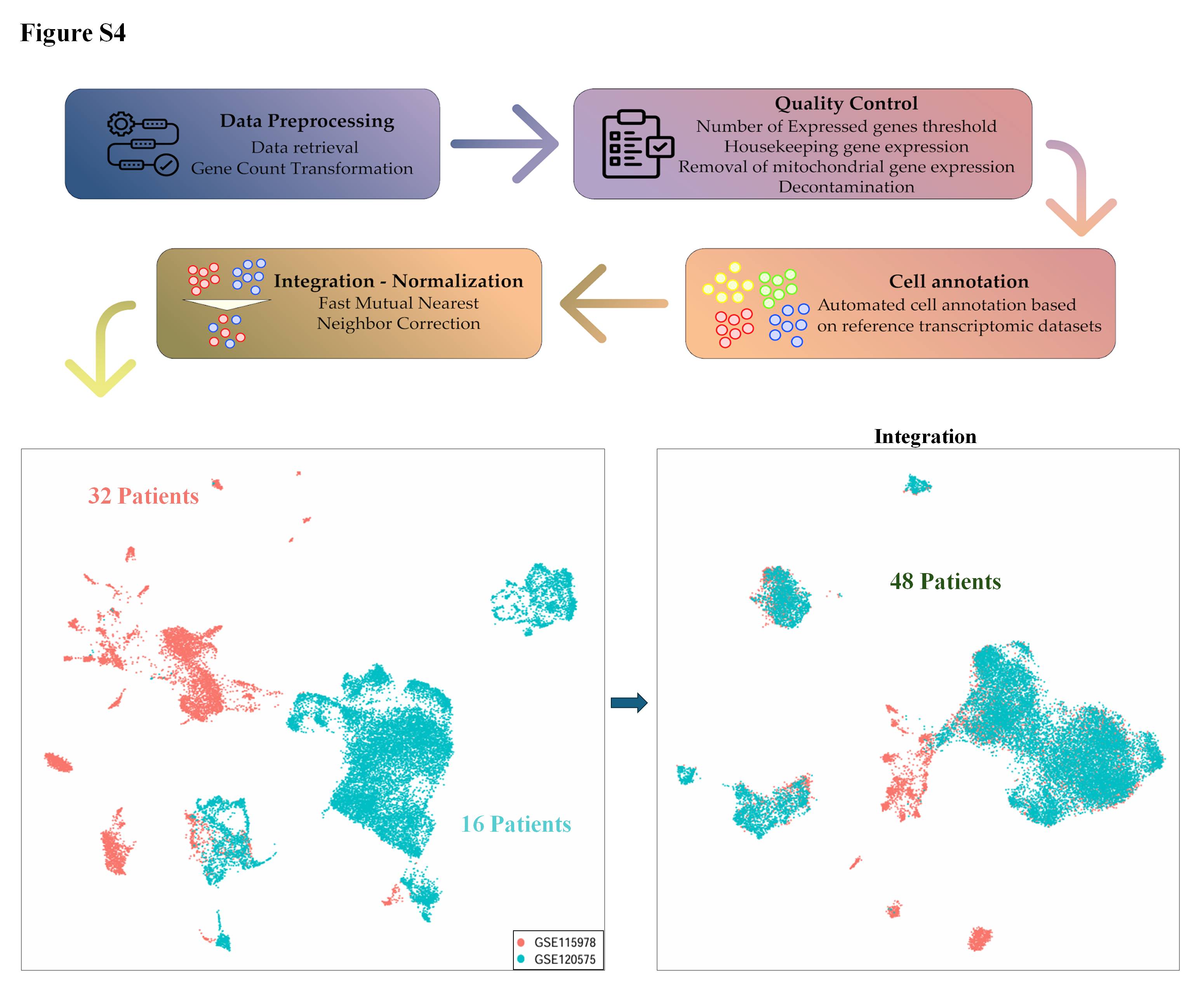

### Suppl Fig 5

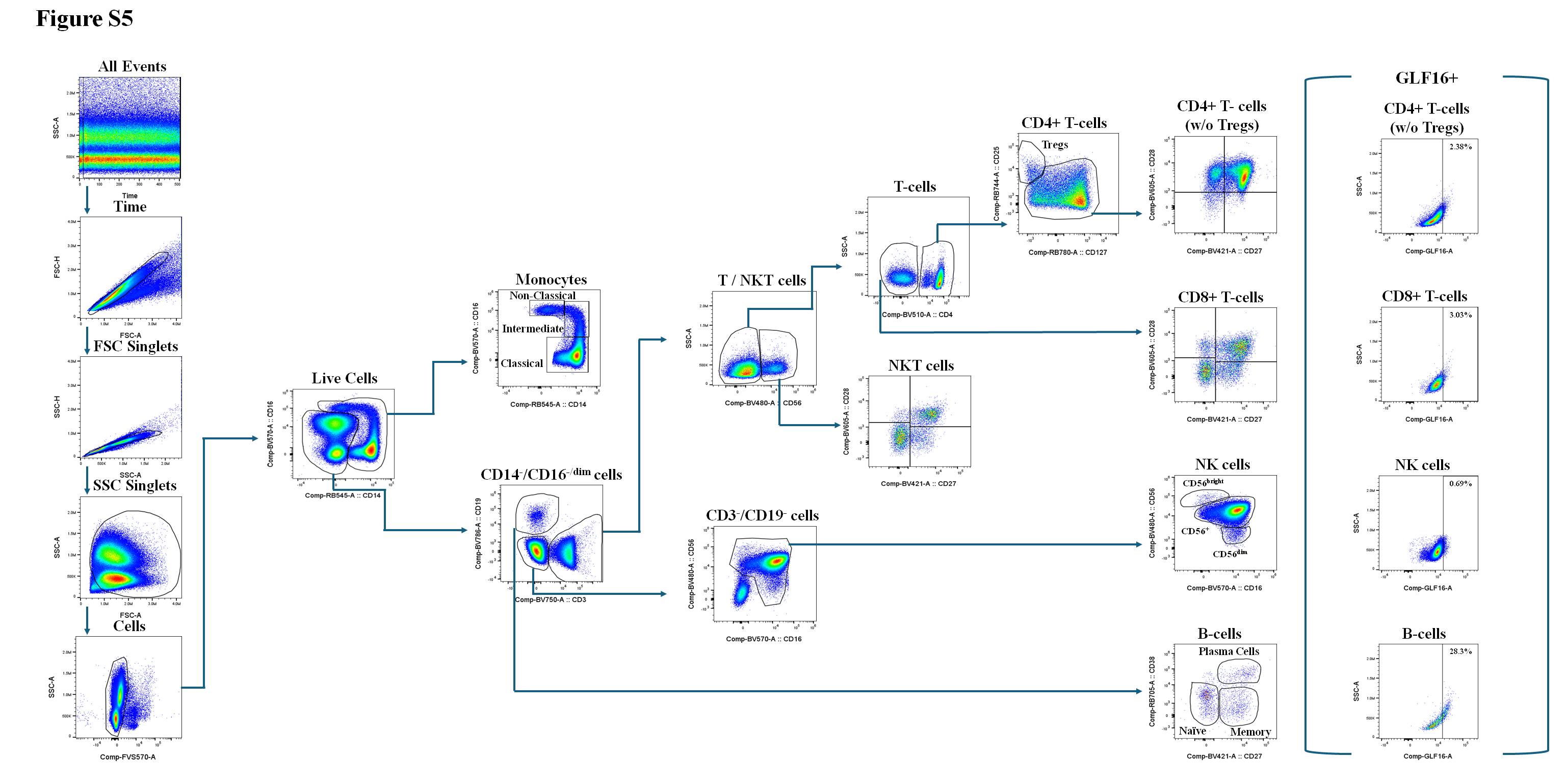

### Suppl Fig 6

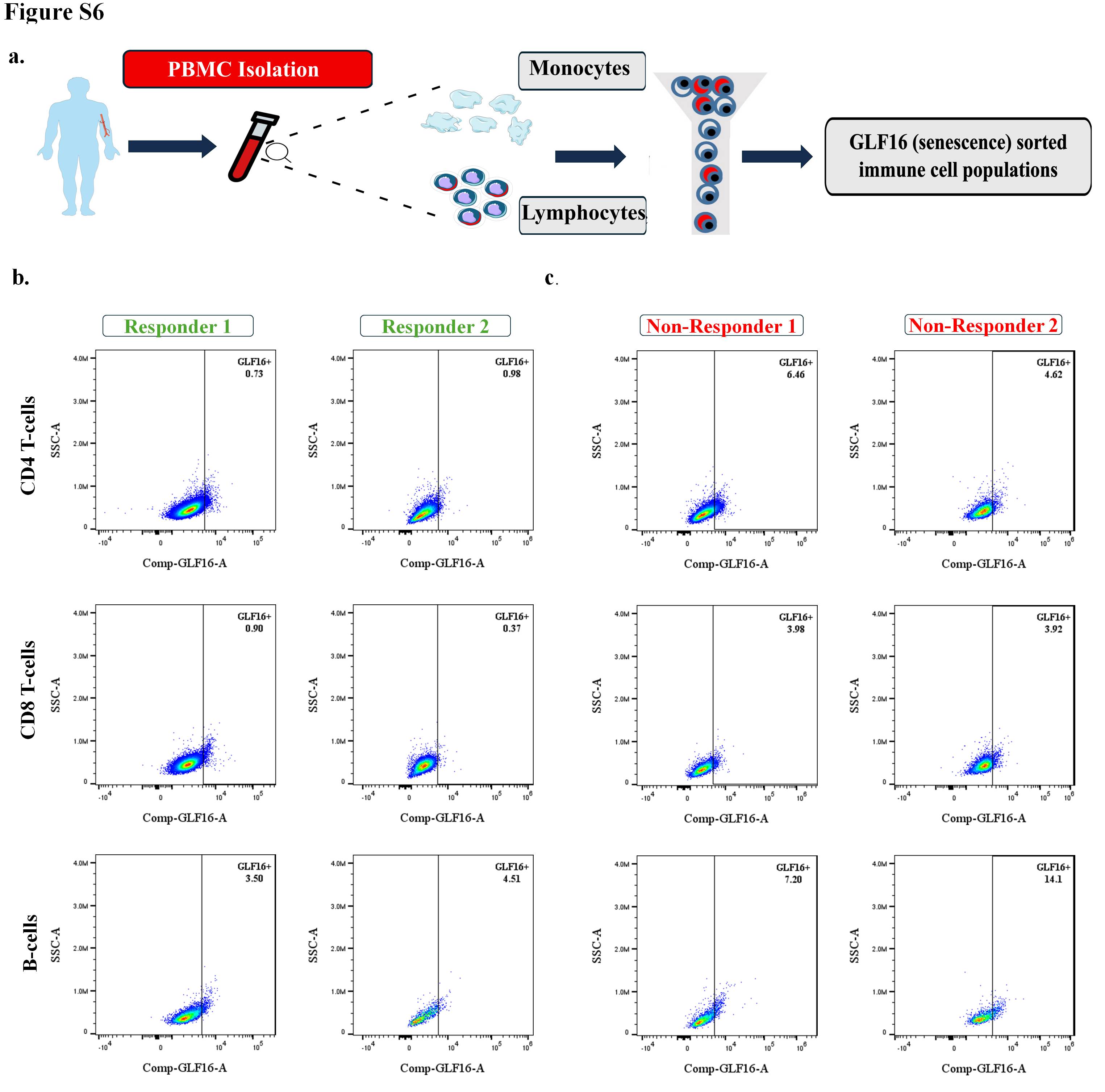

### Suppl Fig 7

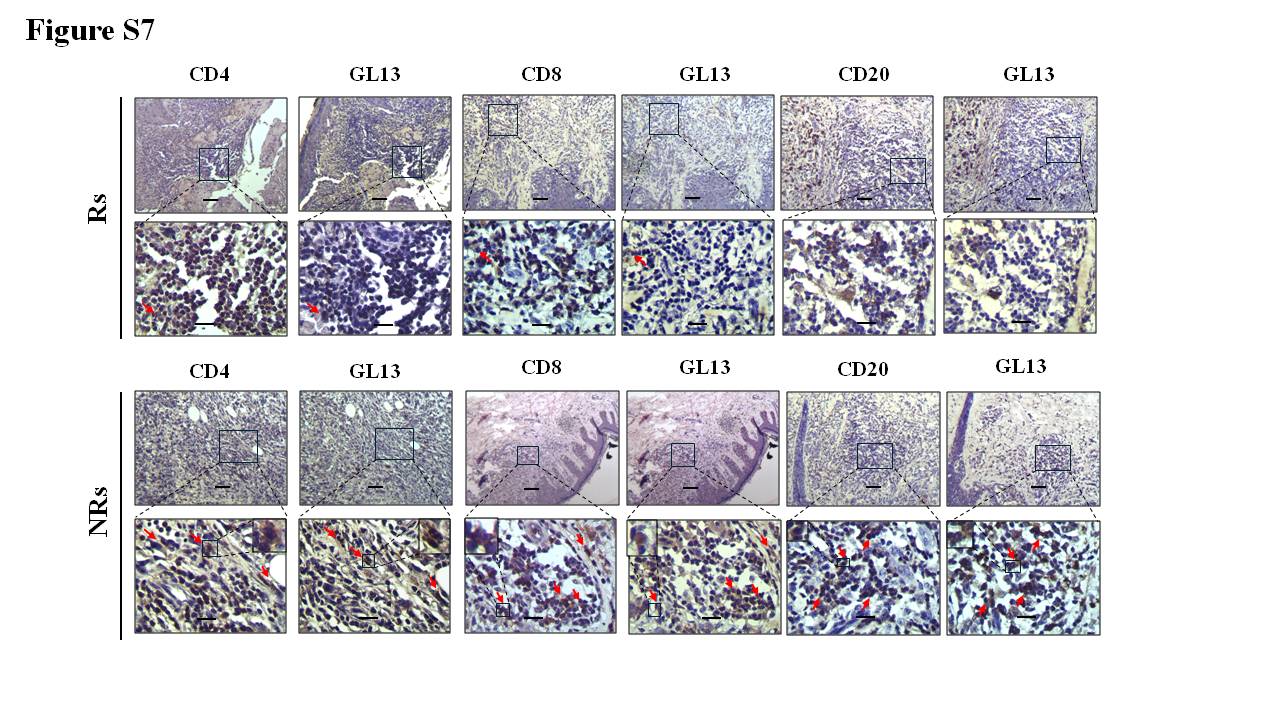

### Suppl Fig 8

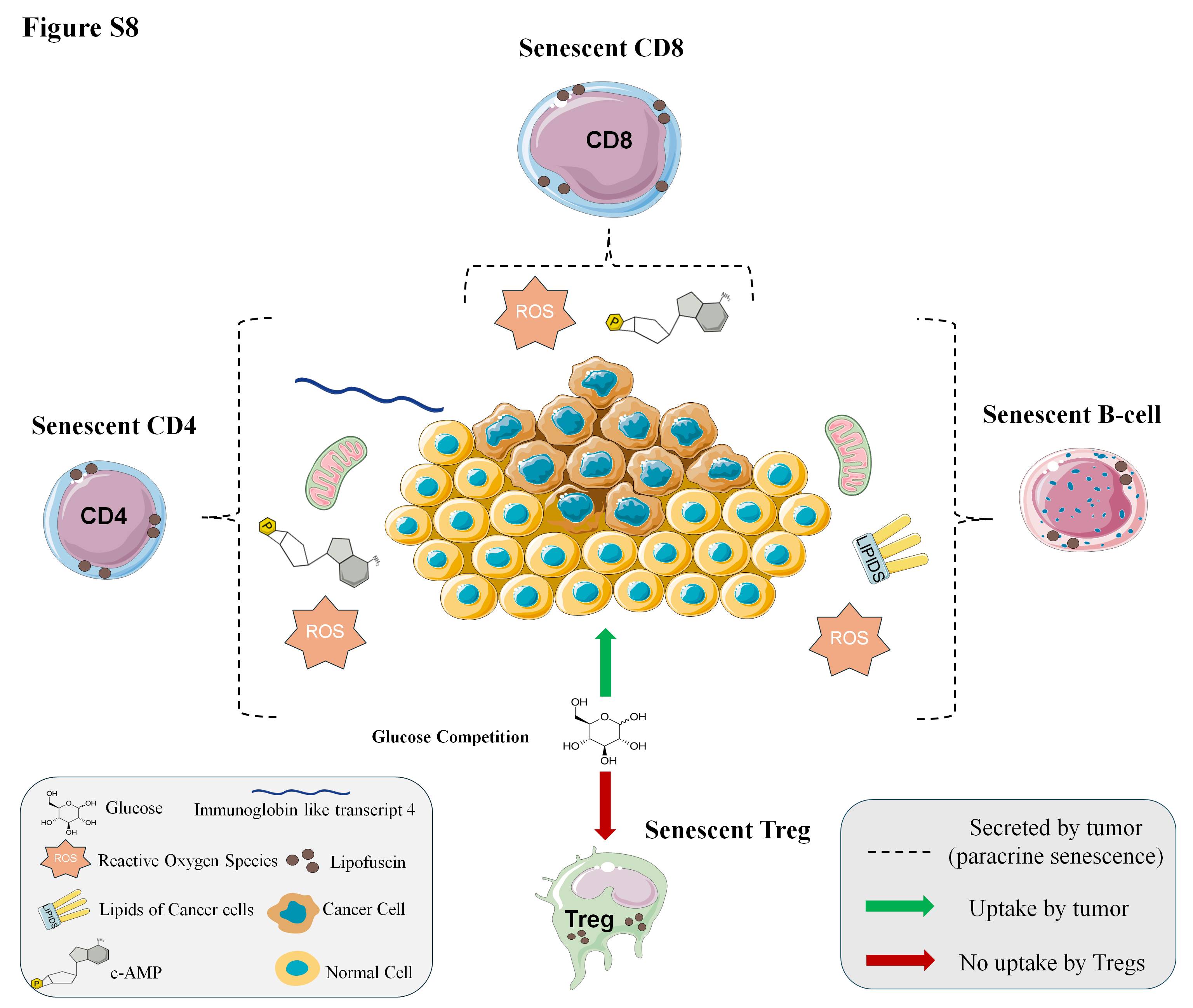
